# Molecular model of a bacterial flagellar motor *in situ* reveals a “parts-list” of protein adaptations to increase torque

**DOI:** 10.1101/2023.09.08.556779

**Authors:** Tina Drobnič, Eli J. Cohen, Tom Calcraft, Mona Alzheimer, Kathrin Froschauer, Sarah Svensson, William H. Hoffmann, Nanki Singh, Sriram G. Garg, Louie Henderson, Trishant R. Umrekar, Andrea Nans, Deborah Ribardo, Francesco Pedaci, Ashley L. Nord, Georg K. A. Hochberg, David R. Hendrixson, Cynthia M. Sharma, Peter B. Rosenthal, Morgan Beeby

**Affiliations:** Department of Life Sciences, Imperial College London, London, SW7 2AZ, UK; Tina Drobnič current affiliation: MRC Laboratory of Molecular Biology, Francis Crick Avenue, Cambridge CB2 0QH, UK; University of Würzburg, Institute of Molecular Infection Biology, Department of Molecular Infection Biology II, Josef-Schneider-Straße 2/D15, 97080 Würzburg, Germany; The Center for Microbes, Development and Health, CAS Key Laboratory of Molecular Virology and Immunology, Institut Pasteur of Shanghai, Chinese Academy of Sciences, Shanghai, China 200031.; Max Planck Institute for Terrestrial Microbiology, Marburg, Germany; Peptone Ltd, 370 Grays Inn Road, London WC1X 8BB, UK; Structural Biology Science Technology Platform, The Francis Crick Institute, London NW1 1AT, UK; Department of Microbiology, University of Texas Southwestern Medical Center, Dallas, TX 75390; Structural Biology of Cells and Viruses Laboratory, The Francis Crick Institute, London NW1 1AT, UK; Centre de Biologie Structurale, Universite de Montpellier, CNRS, INSERM. Montpellier, France

## Abstract

One hurdle to understanding how molecular machines work, and how they evolve, is our inability to see their structures *in situ*. Here we describe a minicell system that enables *in situ* cryogenic electron microscopy imaging and single particle analysis to investigate the structure of an iconic molecular machine, the bacterial flagellar motor, which spins a helical propeller for propulsion. We determine the structure of the high-torque *Campylobacter jejuni* motor *in situ,* including the subnanometre-resolution structure of the periplasmic scaffold, an adaptation essential to high torque. Our structure enables identification of new proteins, and interpretation with molecular models highlights origins of new components, reveals modifications of the conserved motor core, and explain how these structures both template a wider ring of motor proteins, and buttress the motor during swimming reversals. We also acquire insights into universal principles of flagellar torque generation. This approach is broadly applicable to other membrane-residing bacterial molecular machines complexes.

## Introduction

How evolution innovates remains a fundamental question. While innovations in eukaryotes often arise by rewiring existing gene transcriptional networks^1^, examples of the emergence of evolutionary novelty at the molecular scale remain scarce. The few case studies in the emergence of new molecular structures are limited to small protein complexes^2–5^.

Bacterial flagella, helical propellers rotated by a cell-envelope-embedded rotary motor, are examplars of the emergence of molecular novelty^6^ (Fig. 1A). The flagellum, best studied in model organisms *Salmonella enterica* serovar Typhimurium and *Escherichia coli*, is composed of a ring of inner-membrane motor proteins (“stator complexes”) which harness ion flux to rotate a large cytoplasmic rotor ring (the “C-ring”). Torque is transmitted through a chassis (the “MS-ring”) and periplasm-spanning axial driveshaft (the “rod”) to an extracellular propeller structure that generates thrust. Structures of isolated C-ring^7,8^, parts of stator complexes^9,10^, MS-ring attached to the rod^11,12^, and axial structures^13,14^ have recently been determined.

**Figure 1.**
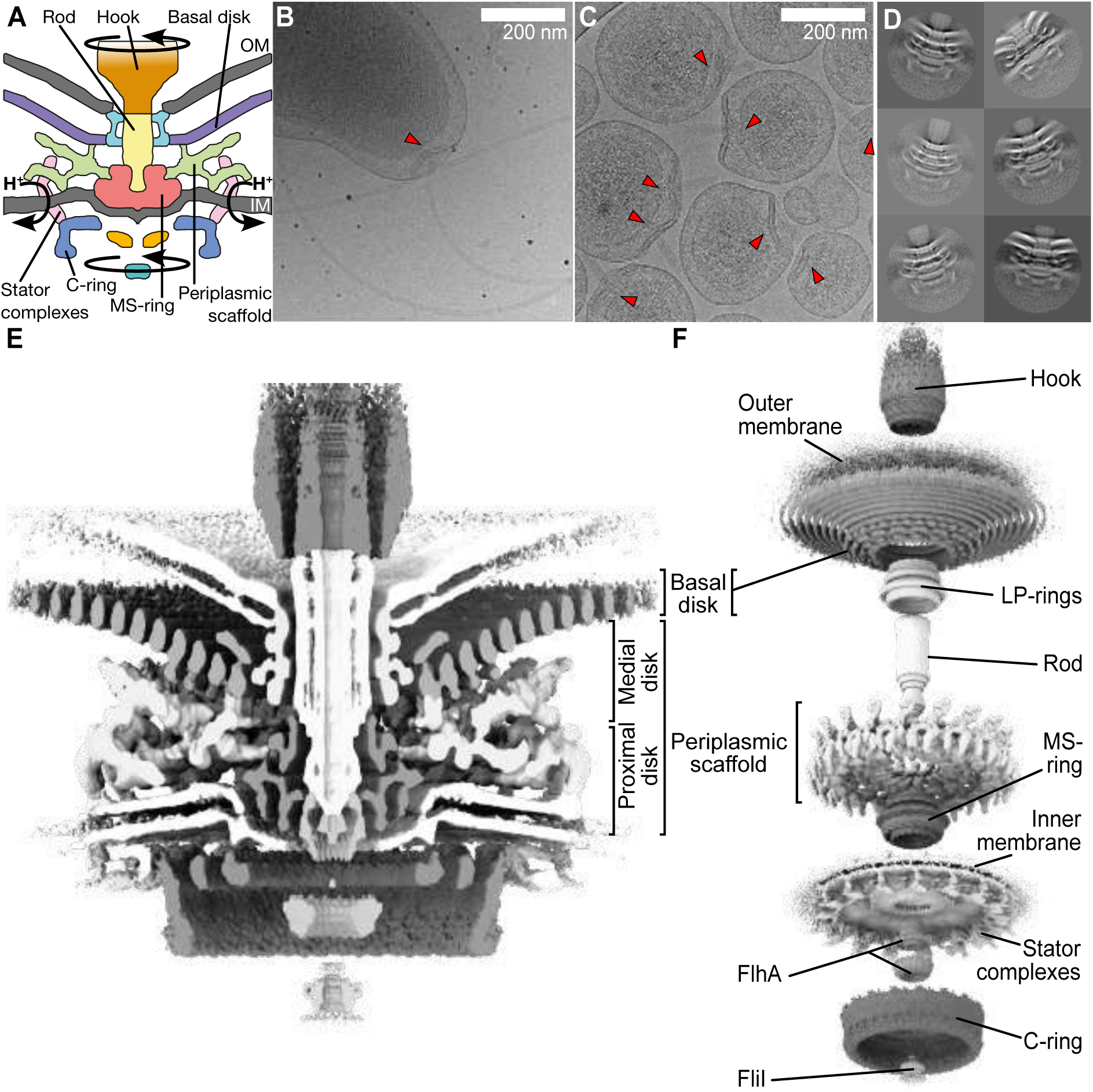
Engineering of homogenous *Campylobacter jejuni* minicells enabled us to determine the *in situ* structure of a flagellar motor by single particle analysis electron cryo-microscopy (A) Schematic of the flagellar motor from *C. jejuni.* Proton flux through the stator complexes drives rotation of the C-ring, MS-ring, rod, and hook/filament. In *C. jejuni* and other Campylobacterota, a basal disk and periplasmic scaffold have evolved that scaffold a wider ring of additional stator complexes thought to be increase motor torque. IM/OM: inner/outer membrane (B) wildtype *C. jejuni* cells typically provide 1 flagellar motor per field of view (arrowhead) as compared to (C) many motors per field of view in our minicell strain (arrowheads), greatly increasing throughput and reducing sample thickness for higher quality electron micrograph acquisition. Note that curvature of minicells is comparable to wildtype cells (D) Periplasmic and cytoplasmic features are evident in single particle analysis 2-D classes of manually picked motors. (E) Cross-section through an isosurface rendering of a C17 whole-motor 3-D reconstruction. (F) Map from (E), segmented and exploded along the z-axis to highlight component structures.

Many flagellar motors have evolved diverse additional proteins associated with greater than the ∼11 stator complexes seen in the model motors^15–17^, likely explaining their associated higher torque. Some of the most complex motors are found in the Campylobacterota, a phylum that includes pathogens *Campylobacter jejuni* and *Helicobacter pylori*^16,18^, which incorporate many additional proteins believed to contribute to three times higher torque generation than *E. coli* and *Salmonella*^17^.

Comparisons of diverse Campylobacterota motors and contextualisation against the *C. jejuni* motor, the most-studied Campylobacterota motor, have suggested a stepwise evolutionary recruitment of proteins^17^. First, inner membrane-associated periplasmic proteins (PflA and PflB, from “Paralysed flagellum” proteins A and B) template assembly of a wider ring of the stator complex motor proteins^16^. This wider ring can exert greater leverage on the correspondingly-wider C-ring to produce the motor’s substantially higher torque, an adaptation likely associated with the highly viscous gut mucous that many Campylobacterota species inhabit^16,18^. Independently, a large (∼100 nm-wide) outer membrane-associated “basal disk” of protein FlgP evolved^16,17,19^ that we recently found is needed for buttressing the motor while wrapping and unwrapping the flagellar filament from the cell body^20^. These structures have been suggested to have initially been independent, and subsequently become co-dependent upon for assembly^17^, but the physical basis for this, and the origins of these new proteins, remains unclear.

Understanding how and why these additional proteins were incorporated into the motor, and how they contribute to function, requires molecular-scale models, but the size, intimate membrane association, and multiple moving parts have hampered structural determination. Cryogenic electron microscopy (cryoEM) subtomogram average structures to between 16 and 18 Å-resolution^11,21^, and structures of purified flagellar motors by single particle analysis cryoEM lacking dynamic components^8,11,12^ do not discern the molecular architecture of these additional structures. Meanwhile advances in imaging membrane proteins *in situ* remain restricted to spherical liposomes rather than native context^22,23^.

To address these challenges, we engineered *C. jejuni* minicells for subnanometre-resolution structure determination of the flagellar motor *in situ.* The quality of our map enabled us for the first time to assemble a molecular model of its evolutionary elaborations. Our model provides key insights into the contributions of new proteins, reveals distant homologies based on structural comparisons, identifies previously unknown components, and contextualises adaptations of pre-existing core machinery to these recruited proteins. We also gain insights into the function of the conserved core of the flagellar motor.

## Results

### Projection imaging of bacterial minicells enables subnanometre resolution *in situ* structure determination

To increase particle number for high-resolution structure determination, we exploited the multiply-flagellated, still-motile minicell phenotype produced by a Δ*flhG* mutation^24^, but removed the flagellum by deleting filament proteins *flaA* and *flaB* for centrifugal purification (Fig. 1B,C). The resulting ∼200 nm-diameter minicells are more homogenous than ∼350 nm-diameter minicells from Enterobacteriaceael minicells such as *E. coli* and *Salmonella*^25^. The polar curvature remains comparable to wildtype cells, meaning motor structures are unperturbed (Fig. 1B,C).

We acquired micrographs of minicells for *in situ* single particle analysis. Initial 2-D classification revealed features including the stator complexes and their periplasmic scaffold, basal disk, C-ring, MS-ring, and rod (Fig. 1D). Classification and refinement applying the established dominant C17 symmetry^16^ yielded a reconstruction to 9.4 Å resolution using 32,790 particles (Fig. 1E, Extended Data Fig. 1). The 17-fold symmetric periplasmic structures exhibited features consistent with discrete proteins, while symmetry-mismatched regions were cylindrical averages of respective structures (Fig. 1E,F). Focused refinement of the periplasmic region improved resolution to 7.9 Å (Fig. 2A; Extended Data Fig. 1D,E,F), sufficient to resolve α-helices and β-sheets (Fig. 2A,B).

**Figure 2.**
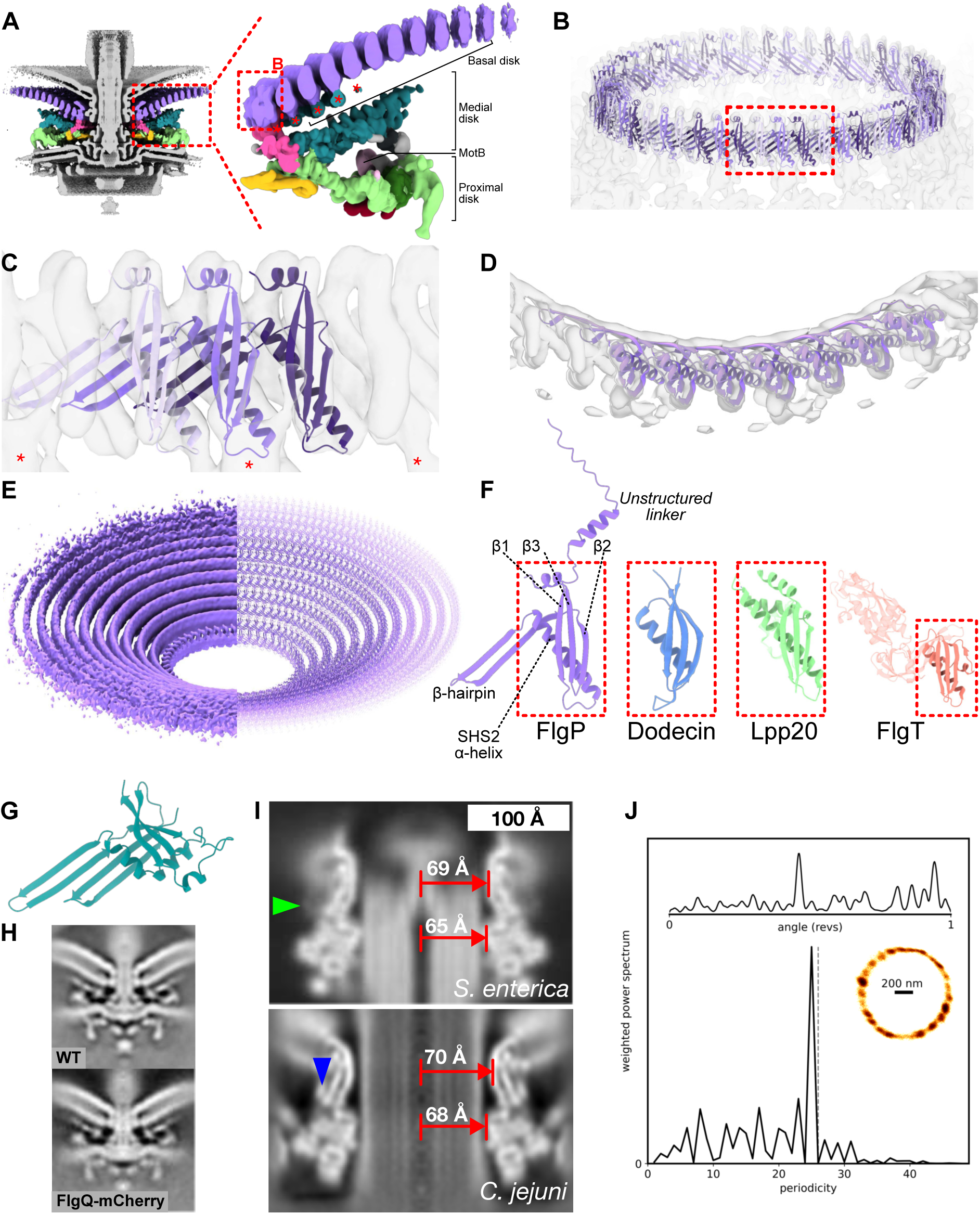
The basal disk is comprised of many concentric rings of FlgP that encircle the LP-rings. (A) Focused C17 refinement of the periplasmic scaffold and inner ring of the basal disk shows that the scaffold attaches to the innermost ring of the basal disk, which itself is a series of concentric rings. Different sections of the periplasmic scaffold and basal disk in colour. (Right) One asymmetric unit of the periplasmic scaffold structure in side view highlighting components. Asterisks denote additional densities attached beneath the first five rings of the basal disk. Note that second and subsequent rings of the basal disk were not part of the focused refinement. (B) The innermost basal disk ring fits 51 FlgP monomers as 17 trimeric protomers (C) Fit of a FlgP trimer into density. Each protomer interacts with the 17-fold symmetric density of the medial disk (asterisks). (D) Top view of a fit of seven FlgP monomers into density map. (E) Density of 10 concentric rings (left) and fitted models (right). (F) FlgP has an unstructured N-terminal linker followed by an SHS2-like fold exemplified by dodecin (PDBID: 1MOG), and shared with Lpp20 of *Helicobacter pylori* (PDBID: 5OK8) and FlgT of *Vibrio alginolyticus* (PDBID: 3W1E). All are OM-associated and FlgT is a flagellar component, indicating shared evolutionary origin. (G) AlphaFold model of FlgQ with signal sequence removed has a double β-hairpin that resembles a two-protomer repeat of FlgP. (H) Tagging FlgQ with mCherry, the resulting motor is indistinguishable from WT. (I) The *Campylobacter jejuni* LP-rings have comparable diameters to the 26-fold symmetric *Salmonella enterica* serovar Typhimurium LP-rings. The density map of the *Salmonella* LP-rings (EMD-12183) was low-pass filtered to 15 Å-resolution and lathed to enable like-for-like comparison with the *C. jejuni* LP-rings. The *Salmonella* L- and P-rings are 69 Å and 65 Å diameter, compared to 70 Å and 68 Å in *C*. *jejuni* Green arrowhead indicates the location of *Salmonella* YecR; blue arrowhead indicates the location of unidentified *C. jejuni* density. (J) Support for the *C. jejuni* LP-rings having comparable stoichiometry to *E. coli* and *Salmonella* from detection of close to 26 steps in flagellar rotation. Top: kernel density estimation of bead position as a function of rotational angle. Bottom: weighted power spectrum of angular position; grey dashed line marks 26 to guide the eye; this example is most consistent with 25 steps. Inset: x,y-position histogram with density represented by darkness of coloration. See Extended Data Fig. 4 for ten other traces.

### The basal disk is composed of concentric rings of FlgP

The basal disk forms a concave cup that pushes the outer membrane away from the inner membrane at increasing distances from the motor (Fig. 2A). Knowing that the disk is composed of FlgP and possibly FlgQ^16,19^, we predicted monomeric and multimeric structures of FlgP, excluding the N-terminal signal sequence and linker^19^ to the outer membrane^20^ using AlphaFold^26^ (Fig. 2A-D). Oligomers laterally associate to form a continuous β-sheet, with interaction of one FlgP with the next two protomers. Bending the arc of these oligomers demonstrated fit of 17 trimeric repeats of 51 protomers into the 51 periodic densities of the inner ring of the basal disk, with a map-model FSC of 9.9 Å at FSC = 0.5 (Fig. 2A-D, Extended Data Fig. 2A, 3A). Each trimeric repeat also featured an additional medial disk-facing density (Fig. 2A,B,C). Although we could not discern the symmetries of rings beyond the 51-fold symmetry of the first, assembly of the disk is entirely reliant upon *flgP*^16^. Subsequent rings all have comparable cross-section densities, making them very unlikely to be composed of a protein other than FlgP. Based on ratios of circumferences, we predict each subsequent ring incorporates 11 additional protomers, explaining the symmetry mismatch with the 17-fold symmetric structures (Fig. 2E).

The FlgP fold is a modified SHS2 domain from InterPro family IPR024952, with a three-stranded β-sheet and an α-helix spanning one face^27^ (Fig. 2F). Structural searches revealed this fold is also present in γ-proteobacterial FlgT, a flagellar component from sodium-driven flagellar motors, and *Helicobacter* Lpp20^28^ (Fig. 2F). FlgP, however, features an additional short C-terminal helix and a long β-hairpin between the α-helix and second β-sheet that extends 35 Å at a ∼42° angle to the vertical axis; it is this β-hairpin that forms the continuous β-sheet in the inner face of each FlgP concentric ring.

FlgQ occurs in an operon with FlgP, is outer membrane-localized, and required for FlgP stability^19^. Its predicted structure resembles a two-protomer FlgP repeat (Fig. 2G). To clarify its location we fused an mCherry tag to FlgQ and determined a subtomogram average structure. Although the mutant was motile, the structure featured no additional densities (Fig. 2H), and we surmise that FlgQ is a low-abundance or irregular component, or an assembly chaperone.

The basal disk is adjacent to the outer face of the P-ring. While ascertaining the symmetry of the L- and P-rings was beyond the resolution of our map, cross-sections through L- and P-ring densities showed comparable radii to *Salmonella*^11,12^ (Fig. 2I). To understand the relationship of the LP-rings to the basal disk, we cylindrically-averaged our map by applying arbitrary high-order symmetry during focused refinement. We then compared these to low-pass filtered structures of similarly cylindrically averaged *Salmonella* LP-rings. The *C. jejuni* L- and P-ring diameters are approximately 103% and 104% those of *Salmonella*, respectively (Fig. 2I), and the lack insertion and deletions, suggested comparable stoichiometries. To experimentally probe the *C. jejuni* LP-ring stoichiometries, we looked for steps in flagellar rotation like the 26 steps imposed by interactions between the asymmetric rod and 26-fold symmetric *Salmonella* and *E. coli* LP-rings^11^. We attached beads to truncated flagella, and upon slowing rotation by de-energizing cells with CCCP, observed 25 to 27 phase-invariant dwell positions (Fig. 2J, Extended Data Fig. 4). Taking these structural and biophysical observations together, we conclude that the *C. jejuni* LP-rings have not expanded to match the symmetry of the basal disk or other novel components, and have a symmetry similar to the 26-fold symmetry of the *Salmonella* LP-rings.

We did not see evidence of YecR, which in *Salmonella* forms a belt around the LP-rings^11^, nor find a homolog in the *C. jejuni* genome, although an unidentified adjacent density hints at an as-yet-unidentified analog (Fig. 2I).

Although our density maps show close association of the P-ring with the basal disk, the basal disk can assemble even when pushed out of axial register with the P-ring by increasing the length of the FlgP N-terminal linker^20^. This indicates that the basal disk does not directly template upon the P-ring, despite their proximity.

### The medial disk is a lattice of PflC with PflD

The medial disk, a lattice between the basal and proximal disks (Fig. 3A), is of unknown composition. An unpublished Tn-seq based infection screen using *C. jejuni* NCTC11168 (Alzheimer, Svensson, Froschauer, Sharma, in preparation) revealed proteins Cj1643 and Cj0892c are required for motility and are polarly-localised, but are not required for flagellar filament assembly, suggesting them as peripheral motor components. Co-immunoprecipitation with proximal disk component PflA in *C. jejuni* 81-176 recovered orthologs CJJ81176_1634 and CJJ81176_0901, respectively (Table S1), which we renamed PflC and PflD (Paralysed flagellum C and D). Subtomogram average structures of deletion mutants (Fig. 3B) revealed loss of the proximal and medial disks upon deletion of *pflC* and loss of a peripheral cage-like structure between the medial and proximal disks upon deletion of *pflD*.

**Figure 3.**
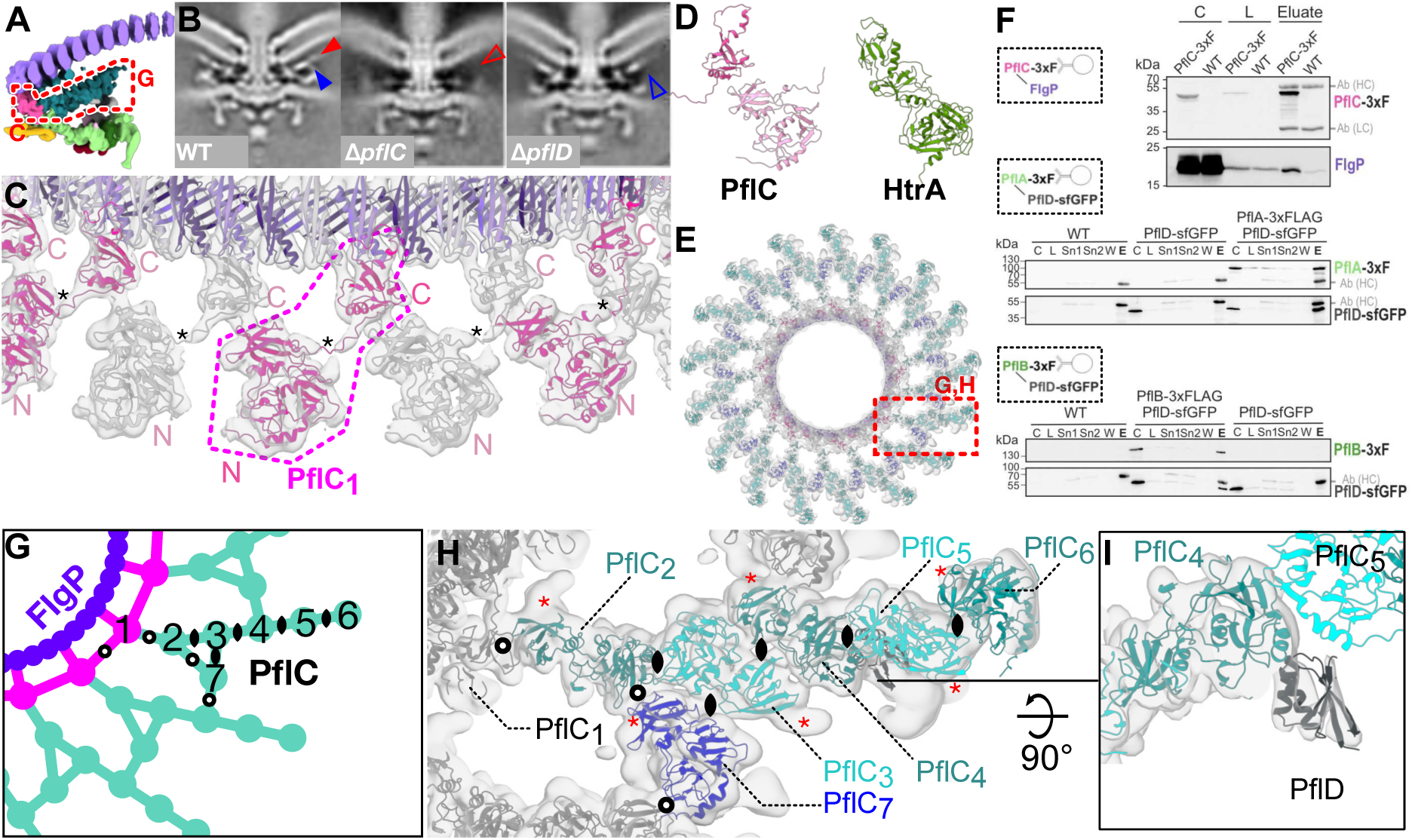
The medial disk is composed of a lattice of previously-unidentified PflC, decorated with PflD and interacts with the basal and proximal disks. (A) The medial disk is situated between the basal disk and proximal disk. (B) Comparing the WT motor structure (from EMD-3150) to a *pflC* deletion reveals abolished assembly of the medial disk (empty red arrowhead; filled red arrowhead on WT structure), while *pflD* deletion abolishes assembly of a post-like density between the medial and proximal disks (empty blue arrowhead; filled blue arrowhead on WT structure). (C) An inner ring of 17 domain-swapped PflC protomers (alternating grey/pink) attach to every third FlgP in the 51-protomer inner ring of the basal disk (purple). A single PflC protomer (magenta outline) features domains from two proteins. Black asterisks denote location of the linker helix between two domains of one protein (D) Predicted structure of PflC highlights common folds and domain architecture with HtrA, a periplasmic protease. (E) Top view of the medial disk viewed from outside the cell. Every asymmetric unit (dashed red box) features seven PflC proteins and one PflD protein as further investigated in panels G and H. PflC_1_ represented in pink as per panel C; PflC_2,4,6_ in teal; PflC_3,5_ in cyan; PflC_7_ in blue. (F) PflC and PflD interact with known flagellar disk structure components: (top) Western blot analysis of coIP experiment of PflC-3xFLAG. As control, untagged wild-type cells (WT) were used. Detected heavy (HC) and light (LC) antibody chains are indicated. C: culture; L: lysate. (middle) Western blot analysis of coIP experiment of PflA-3xFLAG, PflD-sfGFP double tagged strain. As controls, PflD-sfGFP and untagged wild-type (WT) cells were used. Detected heavy (HC) antibody chains are indicated. C: culture; L: lysate; Sn1/2: supernatant 1/2; W: wash; E: eluate. (bottom) Western analysis of coIP experiment of PflB-3xFLAG, PflD-sfGFP double tagged strain. As controls, PflD-sfGFP and untagged wild-type cells were used. Detected heavy (HC) antibody chains are indicated. C: culture; L: lysate; Sn1/2: supernatant 1/2;W: wash; E: eluate. (G) Schematic illustrating the differential oligomerisation of PflC protomers within the medial disk’s lattice. Twofold symmetry axis symbols highlight symmetric dimerization interfaces with axes approximately perpendicular to the plane of the lattice; empty circles represent asymmetric interfaces. Three equivalent twofolds exist: 2:3 and 4:5; 3:4 and 5:6; and 3:7, although the 3:7 interface is substantially warped (H) Molecular model of the PflC lattice fitted in our density map denoting symmetry elements relating monomers enlarged from red box in panel E. Densities adjacent to every Asn239 denoted by red asterisks correspond to an established glycosylation site of a PflC from a closely-related species. Symmetry elements as per panel F. (I) side view depicting PflD sitting beneath PflC_4,5_.

PflC is a 364-residue periplasmic protein featuring N-terminal serine protease and PDZ-like domains (PflC_N_, residues 16-252), a proline-rich linker, and a second PDZ-like domain (PflC_C_, residues 265-364). By rigid-body docking of PflC_N_ and PflC_C_ we located seventeen copies of PflC in the densities projecting from each third FlgP subunit of the first basal disk ring, yielding a map-model FSC of 9.3 Å at FSC = 0.5 (Fig. 3C, Extended Data Fig. 2A, 3B). This domain arrangement resembles trypsin-like HtrA serine proteases^29^ (Fig. 3D), albeit lacking the catalytic residues (Extended Data Fig. 5A,C). PflC_C_ binds FlgP, while PflC_N_ forms the inner band of the medial disk; a density corresponding to the linker links adjacent PflC protomers, revealing that the PflC_C_ domain of one chain assembles with the PflC_N_ domain of another in a daisy-chain of domain-swapped protomers (Fig. 3C). PDZ domains interface with binding partners via a hydrophobic pocket using β-strand addition, but PflC_C_ interactions are unlikely to be mediated by canonical binding^30^ as its ligand-binding groove is oriented away from FlgP, lacks the conserved binding loop^31^, and is predicted to be occupied by residues from its own chain.

The remainder of the medial disk is a lattice of α-helical densities. The resolution was sufficient for us to see that it is composed of six additional PflC_N_ subunits related by diverse dimerization interfaces (Fig. 3E). The PflC_N_ subunits could be docked into our density map with a map-model FSC of 9.7 Å at FSC = 0.5 (Extended Data Fig. 2A, 3C). We refer to the subunits in one asymmetric unit as PflC_2-7_; PflC_2-6_ form radial spokes with a slight twist, while PflC_7_ connects spokes. Validating our assignment of PflC, we noted an additional rodlike density protruding from the location of N239 in all PflCs, which in a closely-related species is glycosylated with an N-linked heptasaccharide^32^. Pulldowns using FLAG-tagged PflC verified the interactions with FlgP and proximal disk components PflA and PflB evident in our map (Fig. 3F).

The PflC lattice consists of diverse oligomerisation interfaces (Fig. 3G,H). There are three types of symmetric dimerisation interfaces with rotational axes perpendicular to the plane of the lattice: one is present in PflC_2_:PflC_3_ and PflC_4_:PflC_5_; one in PflC_3_:PflC_4_ and PflC_5_:PflC_6_; and a third in PflC_3_:PflC_7_, although the latter is substantially distorted. The remaining interfaces (PflC_1_:PflC_1_, PflC_1_:PflC_2_, PflC_2_:PflC_7_, and PflC_7_:PflC_4_) are asymmetric and differ in each case. Based on our findings with PflC_1_, the densities on the underside of the basal disk are likely the C-terminal PDZ domains of PflC_2-7_ (asterisks in Fig. 2A, opaque teal in 3A), although the lack of C17 symmetry renders linker helices invisible.

These promiscuous self-self interfaces predicted that PflC would oligomerise *ex situ*. Curiously, size exclusion chromatography (Extended Data Fig. 5Ei,ii) and mass photometry (Extended Data Fig. 5Eiii) of heterologously expressed PflC indicated a dimer. The *in situ* PflC_N_-PflC_C_ daisy-chaining suggests that the two domains of discrete PflC polypeptides might self-associate, reducing the abundance of the dimer. To test this, we removed PflC_C_. PflC_N_ produced dimers that were more abundant than full-length PflC dimers (Extended Data Fig. 5iv-vi). We speculate that interaction of PflC_N_ and PflC_C_ from the same polypeptide prevents cytoplasmic oligomerization, and binding of PflC_C_ to the basal disk catalyzes assembly of the medial disk only in the context of the assembled motor.

The second candidate medial disk component, PflD, is a 162-residue periplasmic protein that pulls down with PflA of the proximal disk (Table S1). We inspected the peripheral part of the medial disk adjacent to PflC_4N_ which disappeared when we deleted *pflD*, and found that a model of PflD was consistent with this density, despite the lower resolution of this area of our map yielded only a modest mean main chain correlation coefficient of 0.63 (Fig. 3I, Extended Data Fig. 2B).

### PflA spokes, a rim of PflB, and 17 arcs of FliL make the proximal disk

Finally, we sought to interpret our density map of the proximal disk, known to contain PflA and PflB, together with stator complex protein MotB^16^. This region features the extensive short antiparallel α-helical motifs characteristic of the repetitive TPR motifs^33,34^ predicted for proximal disk components PflA and PflB (Fig. 4A). PflA is predicted to form an elongated superhelix consisting of 16 TPR motifs connected to an N-terminal β-sandwich domain by an unstructured linker. Meanwhile PflB is predicted to be α-helical except two five-residue β-strands. We flexibly fitted these structures into our map, with reference to the predicted structure of a PflAB heterodimer (Extended Data Fig. 3D), with 17 radial spokes of PflA positioning a continuous rim of 17 PflBs (Fig. 4B). PflA fitted the map with a map-model FSC of 9.1 Å at FSC = 0.5 (Extended Data Fig. 2A). The resolution of PflB was lower, presumably because it is more peripheral and therefore more flexible, with map-model mean main chain correlation coefficient of 0.71 (Extended Data Fig. 2B).

**Figure 4.**
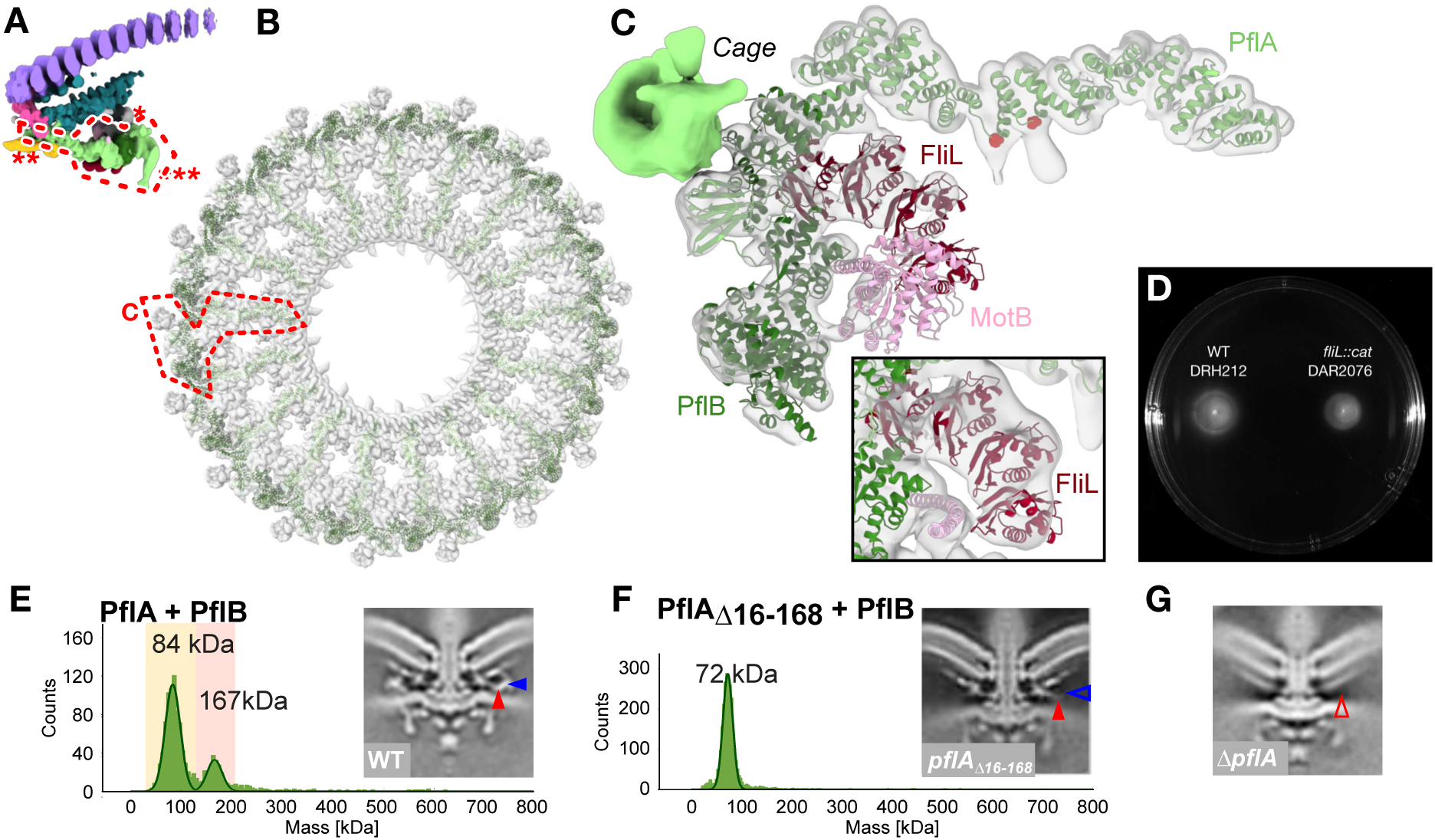
PflA and PflB form a spoke-and-rim structure that scaffolds 17 stator complexes. (A) Location of the proximal disk in the periplasmic scaffold denoted by dashed red line. Asterisks denote unassigned densities: the PflD-adjacent density (*), E-ring (**), and peripheral cage (***) (B) Top view of the 17-fold symmetric proximal disk. Dashed red box denotes the asymmetric unit illustrated in panel C. (C) Every asymmetric unit features one PflA, one PflB, an arc of four FliL, and one stator complex (itself composed of five copies of transmembraneous MotA and two copies of periplasmic MotB). PflA (light green) is positioned radially like spokes, interacting at its N-terminal end with a rim of PflB (dark green) at the outer edge of the scaffold. An arc of FliL (red) and periplasmic domain of MotB (pink, residues 68-247) are also evident at lower confidence. Inset: focus on FliL at lower threshold to demonstrate match of four FliL models into four periodic densities. (D) Deletion of fliL has only a minor effect on motility. A representative motility agar plate stabbed with WT and *fliL::cat* demonstrates that fliL knockout has only a minor effect on motility. (E) Mass photometry measurements confirm the PflAB dimer (red background) forms *in vitro*. Inset: 100 nm x 100 nm cross-section through the subtomogram average density map of the WT motor exhibits PflA_C_ (filled red arrowhead) and PflB densities (filled blue arrowhead). Structure from EMD-3150. (F) Mass photometry shows that deleting the PflA β-sandwich and linker abolishes dimerization with PflB. Inset: 100 nm x 100 nm cross-section through a density map of the subtomogram average of this mutant reveals a vestigial PflA_C_ density (filled red arrowhead) and loss of PflB (empty blue arrowhead), whereas (G) a 100 nm x 100 nm cross-section through a density map of the subtomogram average of a PflA deletion further lacks the vestigial PflA_C_ density (empty red arrowhead) (structure from EMD-3160).

PflAB heterodimer formation is mediated by a the PflA linker binding a TPR-superhelical groove in PflB, with the α-helical PflA spoke pointing toward the motor axis, and the β-sandwich domain wrapping around PflB (Fig. 4C). To verify this model we measured their interaction using mass photometry. PflAB heterodimerised even at nanomolar concentrations (Fig. 4E), while deleting the linker and β-sandwich domain of PflA (residues 16-168) abolished dimer formation (Fig. 4F, Extended Data Fig. 6). A subtomogram average structure of the motor in a PflA_Δ16-168_ mutant confirmed that PflB was unable to assemble into the motor (Fig. 4E), although the C-terminal end of PflA remained evident, unlike a *pflA* deletion mutant^16^ (Fig. 4G). Together with PflAB interaction seen in pulldowns^35^, we conclude that PflAB dimerisation is essential for completion of proximal disk assembly.

Our positioning of PflA is further validated by two additional rodlike densities protruding from the midpoint of each PflA superhelix. Like PflC, PflA is a glycoprotein, with N-linked glycans attached at N458 and N497^36^, and these residues are situated at the base of the likely N-linked heptasaccharide densities (Fig. 4C, red atoms).

An arc of density partially encircling the periplasmic MotB linker had similar radius and location as the tertiary structures of complete circles of FliL in other motors^37,38^, and we found that a curved tetrameric homology model of FliL fitted into this arc with a mean main chain correlation coefficient of 0.61 (Fig. 4D, Extended Data Fig. 2B). Co-immunoprecipitation assays confirmed that FliL is found in pulldowns of PflA and PflB (Table S1), suggesting that the FliL arc is augmented by PflB and PflA to scaffold MotB and explains why PflA and PflB are both required for the high occupancy or static anchoring of stator complexes into the *C. jejuni* motor^16^. Indeed, we found that deletion of *fliL* in *C. jejuni* had only a modest impact upon motility (Fig. 4D) in contrast to *H. pylori* where FliL is essential, reinforcing that FliL’s role is partially fulfilled by PflA and PflB in *C. jejuni*. The presence of the stator complexes in the *C. jejuni* structure, in contrast to their absence in *Salmonella,* indicates either high occupancy or static anchoring, possibly mediated by their interactions with PflA, PflB, FliL, and potential cytoplasmic proteins. Although our previous work and the topology of MotB make its location unambiguous, the resolution of this region was low, meaning we could only crudely position a MotB model.

We could not assign proteins to three remaining densities in the scaffold: the so-called E-ring that spaces the MS-ring from PflA, a cage previously observed in *H. pylori*^37^ on the periphery of the PflB rim that extends through the membrane to wrap around the stator complexes, and a small density adjacent to PflD (Fig. 4A, opaque regions, single, double, and triple asterisks respectively).

### Conserved motor components have adapted to a high torque role

To better understand the flagellar torque-generation machinery we focused on the stator complexes, C-ring, and MS-ring. By symmetry expanding and classifying stator complexes from our whole-motor map, we observed a pentameric structure directly beneath the periplasmic peptidoglycan-binding domain of MotB that contacts the C-ring (Fig. 5A,B,C). In cross-section, the dimensions and shape of this density match that of purified *C. jejuni* stator complex membrane component MotA_5_^10^ (Fig. 5C). The consistent rotational register of this pentameric density even after symmetry expansion and classification, with a pentameric corner pointing toward the C-ring, indicates that stator complexes are most frequently in this rotational register.

**Figure 5.**
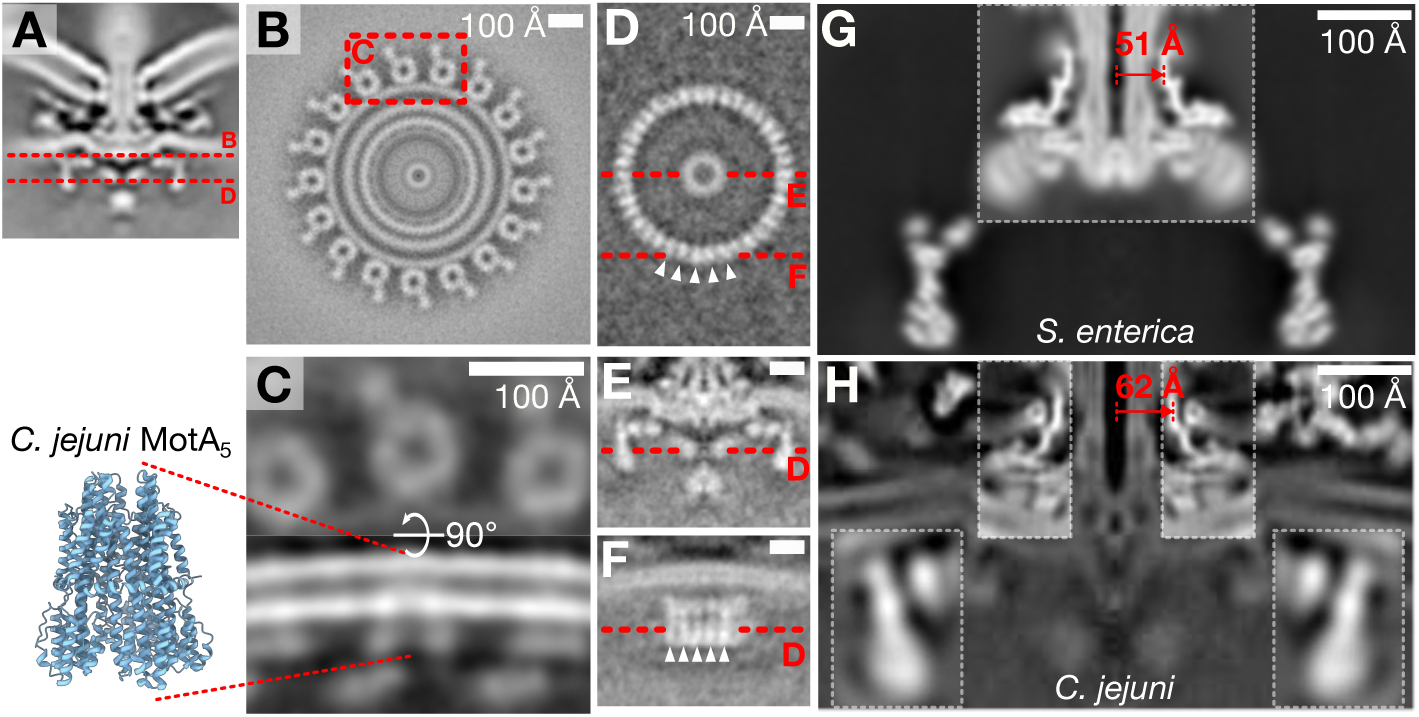
Wider rings of additional stator complexes are incorporated in the *Campylobacter jejuni* periplasmic scaffold while the rotor components are correspondingly wider. (A) Cross-section the wildtype *C. jejuni* motor depicting locations of stator and rotor cross-sections illustrated in panels B and D. (B) Cross-section through the whole-motor map just beneath the outer membrane shows 17 circular densities at the expected location of MotA. (C) A focused refinement of the stator complexes reveals pentameric densities, that (C) In cross-section have the distinctive thimble-like shape of a MotA pentamer, from PDB 6ykm. (D) Cross-section through the *C. jejuni* C-ring showing 38-fold periodic structure. Arrowheads highlight 5 of the 38 puncta. Labels (E) and (F) denote cross-sections depicted in respective panels. (E) Cross-section through the centre of the *C. jejuni* C-ring; (F) cross-section through the edge of the C-ring showing post-like densities corresponding to the periodicity shown in panel (D) (arrowheads) as have been reported in the *Salmonella* C-ring. (G) Cross-section through a composite map of the *Salmonella enterica* serovar Typhimurium MS-ring (from EMD-12195) and C-ring (from EMD-42439) rotor components depicting the 51 Å-radius MS-ring β-collar. Both maps were low-pass filtered to 15 Å-resolution and lathed around their rotational axis to allow like-for-like comparison with (H) cross-section through a composite map of the whole-motor *C. jejuni* map, with superimposed cross-sections through focused, lathed *C. jejuni* MS-ring and C-ring maps, highlighting the wider 62 Å-radius β-collar.

We wondered how the *C. jejuni* rotor components, i.e., the C-ring, MS-ring, and rod, have adapted to interact with a wider ring of stator complexes. The *C. jejuni* C-ring is wider than that of *Salmonella*^16^; to ascertain its stoichiometry, we determined its structure by subtomogram averaging. To remove the strong 17-fold signal of MotA, we used a motile *C. jejuni* mutant whose stator complexes have lower occupancy than WT motors. Our subtomogram average revealed a 38-fold periodic structure whose vertical post-like architecture resembles that of the near-atomic-resolution *Salmonella* C-ring^8^ (Fig. 5D,E,F).

C-ring diameter is reliant upon templating on the C-terminus of the MS-ring protein FliF^8,39^. To examine possible mechanisms behind templating a wider C-ring, we compared the *C. jejuni* MS-ring to that from *Salmonella.* The distinctive FliF β-collar has a radius of approximately 51 Å (Fig. 5G). Focused refinement of the *C. jejuni* MS-ring with imposed C38 symmetry expected from 1:1 stoichiometry of FliF and FliG^39,40^ demonstrated a 62 Å-radius FliF β-collar (Fig. 5H). We could not resolve azimuthal features corresponding to discrete FliF subunits, but imposition of other arbitrary high-order symmetries did not alter the radius of FliF’s β-collar. We conclude that the wider C-ring is achieved by *C. jejuni* assembling a wider MS-ring to template the wider C-ring.

## Discussion

This work provides a near-complete inventory of the proteins incorporated into the *C. jejuni* motor during evolution of higher torque output, adaptations of pre-existing components (Fig. 6, Table S2), and insights into universal principles of flagellar rotation.

**Figure 6.**
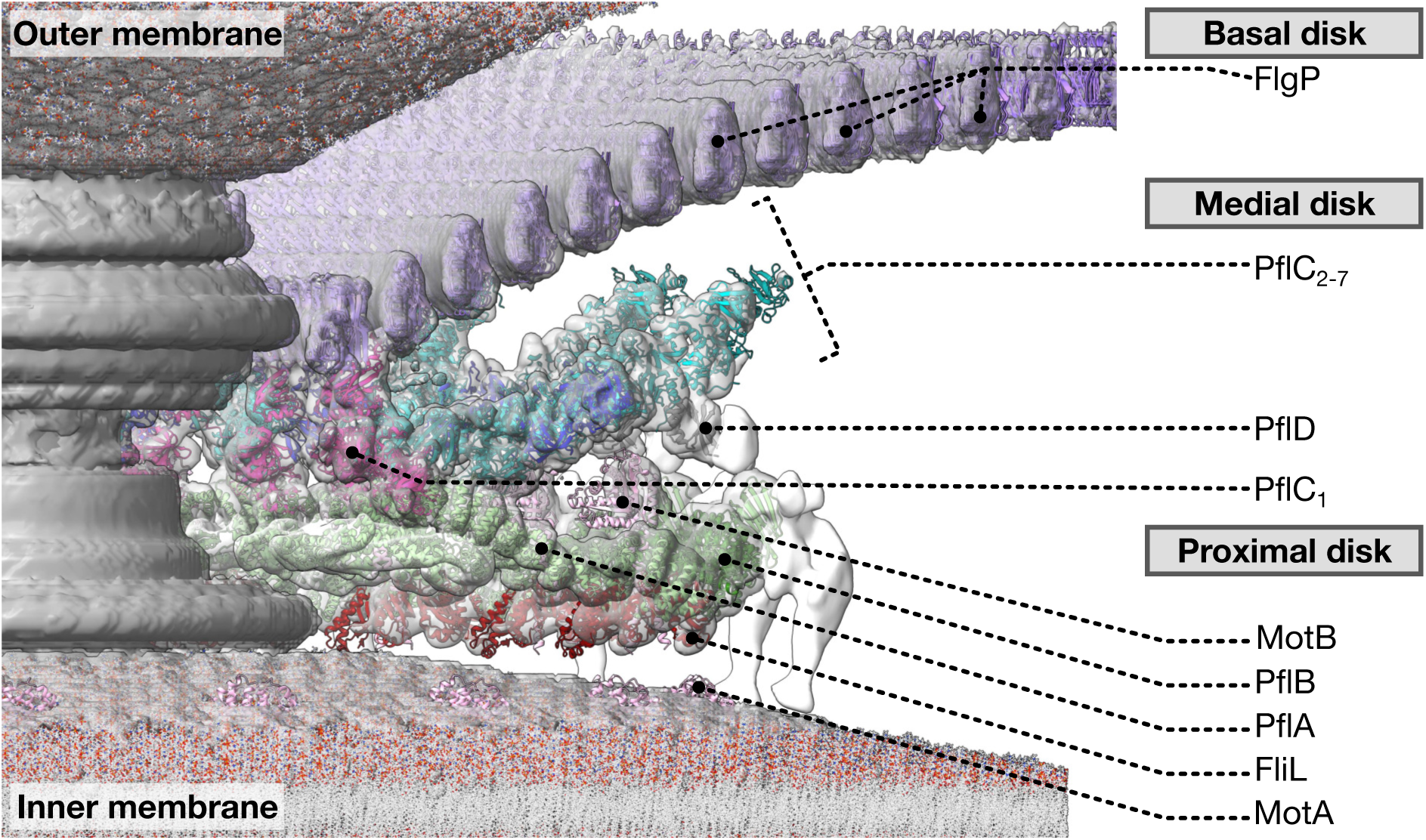
A “parts-list” of protein adaptations to increase torque by scaffolding a wider ring of additional stator complexes, thus exerting greater leverage on the axial flagellum. A partial-cutaway schematic of the structure of the *Campylobacter jejuni* flagellar motor contextualising protein components modelled in this study. The basal disk is formed of FlgP; the medial disk of PflC and PflD; the proximal disk of PflA, PflB, and FliL together with stator complex components MotA and MotB. Asterisks on transparent density denote unassigned densities: a PflD-adjacent density (*), E-ring (**), and peripheral cage (***).

Our structures explain how the *C. jejuni* motor produces approximately three times the torque of the *E. coli* and *Salmonella* motors. The spoke-and-rim scaffold of PflA and PflB, supported by PflC and PlflD, positions stator complexes in a wider ring; these not only each exert greater torque due to a greater lever length, but the torque from additional stator complexes is additive^16,41^. This scaffold facilitates the high occupancy or static positioning of the *C. jejuni* stator complexes^16^; the radius of the MS-ring is correspondingly larger than that of *Salmonella* to template a wider C-ring to maintain contact with the larger stator ring. PflA and PflB are composed of arrays of TPR motifs, widespread building blocks of structural scaffolds. The widespread nature of TPR motifs would have made them available for co-option to form PflA or PflB. Intriguingly, parts of the Tol/Pal system homologous to other flagellar proteins^6^ and the Tol/Pal component YbgF features TPR motifs^42^, suggesting a scenario in which PflA and PflB originated from another Tol/Pal co-option.

*C. jejuni*, the proximal disk is linked to the basal disk by the medial disk, while other species lack a medial disk and have separate proximal and basal disks. Our structures provide insight into the significance and mechanism of the relatively recent evolution of the medial disk^16^. We recently showed that the basal disk has a distinct function in buttressing the motor when unwrapping flagellar filaments from the cell body^20^. Here we identify PflC as the mediator of the extensive connections between the proximal and basal disks^20^ (Fig. 3, 4). PflC’s shared domain organisation and structure with HtrA-family enzymes suggests that it originated by co-option of an HtrA family protein^43^ (Extended Data Fig. 5A,C). Indeed, HtrA family proteins are periplasmic, meaning that they already co-localised with the flagellar motor. Curiously, HtrA also forms higher-order oligomers^29^, suggesting that this oligomerisation tendency was a pre-existing property easily exapted for its role as a structural scaffold. The proline-rich linker between PflC_N_ and PflC_C_ is evidently important for the head-to-tail oligomerisation of PflC, because species such as *Wolinella* and *Helicobacter,* which lack a medial disk, have PflC sequences without this linker.

The basal disk is comparatively more ancient. While the basal disk protein FlgP comes from a broad protein family, ring oligomerisation in the family correlates with the subset of sequences that feature a β-hairpin insertion between the α-helix and second β-sheet, as seen with the outer membrane-associated rings formed by FlgT, whose SHS2 domain features this insertion^44^. Curiously, the basal disk from *Wolinella* has been proposed to form a spiral, not concentric rings^45^, raising the possibility that FlgP forms spirals in the absence of a medial disk.

Our study sheds light on the universal mechanism of flagellar rotation. We discerned a consistent rotational register of MotA, with vertices oriented toward the C-ring, and focused classification of individual stator complexes failed to resolve classes with other rotational registers. This is inconsistent with the homogenous circle predicted for averaged structures given the continuum of rotational registers expected from a high duty ratio of the stator complexes with FliG and the symmetry mismatch between the 17 stator complexes and 38-fold symmetric C-ring. Predictions of stator complexes having a high duty ratio assumed an elastic linker from peptidoglycan to MotA as in *Salmonella*^46^, although this does not hold for *C. jejuni* in which stator complexes are rigidly encased in their proteinaceous cage. Our structure is therefore consistent with recent results suggesting that the stator complexes having a low duty ratio^47^, and MotA’s rotational register is dictated by an energy minimum of rotation around MotB instead of against FliG.

While our projection imaging approach achieves substantially higher resolution than previous studies, enabling molecular interpretation and validation, our approach is not without limitations. Our study focused on the 17-fold rotationally symmetric periplasmic scaffold, but flagellar motors feature diverse symmetry-mismatched subcomponents. The approach can in principle identify arbitrary symmetries, but we did not resolve symmetries for other substructures such as the C-ring, likely due to weak signal due to their size, heterogeneity and excessive noise from the *in situ* context. (Although useful information on these structures is available from their cylindrical averages.) Different data acquisition schemes or mutants are needed to understand how to extract this signal. Nevertheless, using minicells to study polar machinery has promise for substantial increases in resolution in other systems, including Tad pili^48^ in *Caulobacter crescentus* minicells^49^, type II secretion systems^50^ in *Pseudomonas* minicells^51^, chemoreceptors in *C. jejuni*^52^, and the flagellar motors of diverse other polar flagellates^53,54^.

Our approach achieved sufficient resolution of a bacterial molecular machine *in situ* for molecular interpretation. Results provide invaluable information on the mechanisms of torque generation and possible mechanisms of evolution, and will provide a foundation for further comparative studies to probe how novel interfaces arose during protein recruitment. Our approach confirms that imaging protein machines *in situ* can provide subnanometre resolution, and lays a foundation for future studies expanding to other symmetry elements and structures.

## Acknowledgements

We thank Paul Simpson in the Imperial College London Electron Microscopy Centre and the Francis Crick Institute Structural Biology science technology platform for electron microscopy assistance. This work was supported by Medical Research Council grant MR/V000799/1 to EJC and MB, Human Frontier Science Program grant RGP0028/2021-HOCHBERG to MB and GH, NIH grant R01AI065539 to DRH, BBSRC doctoral training grant BB/M011178/1 and a short-term DAAD grant to TD. PBR is supported by the Francis Crick Institute, which receives its core funding from Cancer Research UK (CC2106), the UK Medical Research Council (CC2106), and the Wellcome Trust (CC2106). CMS is supported by research grants Sh580/7-2 and Sh580/8-2 in the framework of the DFG (Deutsche Forschungsgemeinschaft) priority program SPP2002 (Small proteins in prokaryotes, an unexplored world), the DFG Research Training Group GRK2157 “3D-Infect”, and a “CampyRNA” junior consortium grant within the 2nd call of Infect-ERA (ERA-NET; www.infect-era.eu)/Bundesministerium für Bildung und Forschung (BMBF; www.bmbf.de). ALN was supported by two French National Research Agency (ANR) grants (PHY-BABIFO ANR-22-CE30-0034 and PHYBION ANR-23-ERCB-0005-01). The CBS is a member of the France-BioImaging (FBI) and the French Infrastructure for Integrated Structural Biology (FRISBI), two national infrastructures supported by the ANR (ANR-10-INBS-04-01 and ANR-10-INBS-05, respectively).

For the purpose of open access, the author has applied a Creative Commons Attribution (CC BY) license to any Author Accepted Manuscript version arising.

## Author Contributions

TD: sample prep, image processing, analysis, molecular modelling, mass photometry, wrote paper; EJC: strain construction, sample prep, image processing, analysis; TC: image processing, manuscript and figure preparation; NS: image processing; TU: sample prep, data acquisition, image processing; AN: data acquisition, image processing; DR: flgQ-mCherry strain construction, pulldowns, fliL knockout; SS, KF, MA, CS: PflC and PflD identification; WHH, FP, ALN: experimental design, data acquisition, data analysis, funding acquisition; LDH: PflD subtomogram averaging; GH, SGG: Mass photometry data acquisition supervision and analysis, funding acquisition; DRH: strain construction, pulldowns, funding acquisition; PR: image processing, manuscript and figure preparation, funding acquisition; MB: conceptualization, funding acquisition, supervision, wrote paper

## Methods

### Lead contact

Further information and requests for resources and reagents should be directed to and will be fulfilled by the lead contact, Morgan Beeby

### Materials availability

Plasmids and strains generated in this study (see Table S3) are available on request from the lead author.

### Data and code availability

Cryo-EM maps have been deposited at the Electron Microscopy Data Bank (EMDB) and are publicly available as of the date of publication. The whole-motor map has been deposited with accession code EMD-16723 together with the original micrographs deposited to the EMPIAR repository with public accession code EMPIAR-11580 (DOI: 10.6019/EMPIAR-10016). The refined periplasmic scaffold map has been deposited in the EMDB with accession code EMD-16724. Subtomogram average maps were deposited as follows: Δ*pflC* - EMD-17415; Δ*pflD* - EMD-17416; *pflA*Δ16-168 - EMD-17417; FlgQ-mCherry - EMD-17419.

### Bacterial strains and culture conditions

*C. jejuni* 81-176 or NCTC11168 were cultured from frozen stocks on Mueller-Hinton (MH) agar (1.5% w/v) supplemented with trimethoprim (10 μg/mL) (MHT) for 1-2 days at 37 C under microaerophilic conditions (5% O2, 10% CO2, 85% N2) in a in a Heracell 150i trigas incubator (Thermo Fisher Scientific). Additional antibiotics were added to the agar medium when required: kanamycin (Km) at 50 μg/mL, streptomycin (Sm) at 2 mg/mL. All 81-176 mutants were constructed in DRH212^55^, a streptomycin resistant derivative of *C. jejuni* 81-176, which is the reference wild-type strain in this work unless otherwise stated. When working with *E. coli*, cultures were grown at 37 C on Luria-Bertani (LB) agar plates (1.5% w/v) or in LB medium with agitation, both supplemented with carbenicillin at 100 μg/ml. Please see Table S3 for details of strains and plasmids used in this study.

## Method details

### Strain construction

The minicell (Δ*flhG* Δ*flaAB*) and PflA truncation (pflA_Δ18-168_) strains were constructed as described previously^55,56^. Briefly, *aphA-rpsL^WT^* cassettes flanked by ∼500 bp overhangs with homology to the targeted chromosomal loci and ecoRI sites at the 5’ and 3’ termini were synthesised by “splicing by overlap extension” PCR (SOE PCR). Linear DNA fragments were methylated at their ecoRI sites with ecoRI methyltransferase (New England Biolabs) and transformed into *C. jejuni* using the biphasic method^57^. Transformants were selected for on MH agar supplemented with 50 µg/mL kanamycin. Replacement of the *aphA-rpsL^WT^* with the desired mutation was achieved using the same method, but with transformants being selected for on MH agar supplemented with 2 mg/mL streptomycin sulfate. Kanamycin-sensitive, streptomycin-resistant transformants were single-colony purified and checked by Sanger sequencing (Source Biosciences UK). For the minicell background, in-frame deletion of *flhG* leaves the first and last 20 codons intact, while the Δ*flaAB* allele spans from 20 base pairs upstream of the *flaA* translational start site to codon 548 of *flaB*.

To construct the *C. jejuni fliL* mutant, we made a *cat* insertional knockout and confirmed absence of polar effects. To preserve expression of the essential *acpS* gene downstream of fliL, we constructed a *fliL* mutant that disrupted *fliL* with an antibiotic-resistance cassette containing an intact flaA promoter positioned to maintain expression of acpS. First, the *fliL* locus from *C. jejuni* 81-176 was PCR amplified with a HpaI site engineered within the *fliL* coding sequence. This fragment was then cloned into the BamHI site of pUC19 to create pDAR1712. The flaA promoter and start codon were PCR amplified from *C. jejuni* 81-176 and cloned into the XbaI and BamHI sites of pUC19 to create pDAR2039. The chloramphenicol-resistance cassette containing cat was digested as a PstI fragment from pRY109 and cloned into PstI-digested pDAR2039 to create pDAR2045. The *cat-flaA* promoter was then digested from pDAR2045 as a EcoRI-BamHI fragment, treated with T4 DNA polymerase to create blunt ends and cloned into the HpaI site of pDAR1712 to create pDAR2072. pDAR2072 was verified to contain the *cat-flaA* promoter in the correct orientation to maintain expression of *acpS*. DRH212 was then electroporated with pDAR2072 and chloramphenicol-resistance transformants were recovered. Colony PCR verified creation of a *fliL* mutant (DAR2076).

Deletion mutants of *C. jejuni* NCTC11168 were constructed by double-crossover homologous recombination with an antibiotic resistance cassette to remove most of the coding sequence using overlap PCR products. As an example, deletion of *cj1643* (*pflC*) is described. First, ∼500 bp upstream of the *cj1643* start codon was amplified using CSO-3359/3360 and ∼500 bp downstream of the *cj1643* stop codon was amplified with CSO-3361/-3362 from genomic DNA (gDNA) of the wild-type strain (CSS-0032). A non-polar kanamycin resistance cassette (*aphA-3*, Kan^R^) (Skouloubris et al., 1998) was amplified from pGG1^58^ with primers HPK1/HPK2. To fuse the up- and downstream regions of *cj1643* with the resistance cassette, the three fragments were mixed and subjected to overlap-extension PCR with CSO-3359/3362. PCR products were electroporated into the WT strain as previously described^59^. The final deletion strain (CSS-4087; NCTC11168 Δ*cj1643*) was verified by colony PCR with CSO-3363/HPK2. Deletion of *cj0892c* (*pflD*) in C. jejuni strain NCTC11168 was generated in a similar fashion: *cj0892c*::*aphA-3* (CSS-4081; NCTC11168 Δ*cj0892c*).

To fuse *sfgfp* fusion to the penultimate codon of *cj0892c* (*pflD*), its coding sequence was first amplified with CSO-3611/3612, digested with *Nse*I/*Cla*I, and inserted into similarly-digested pSE59.1^60^; amplified with CSO-0347/CSO-0760) to generate pSSv106.5, where *cj0892c* transcription is driven from the *metK* promoter. The plasmid was verified by colony PCR with CSO-0644/3270 and sequencing with CSO-0759. Next, *sfgfp* was amplified from its second codon from pXG10-SF^61^ with CSO-3279/3717, digested with *Cla*I, and ligated to pSSv106.5 (amplified with CSO-3766/0347 and also digested with *Cla*I). This generated pSSv114.1, which was verified by colony PCR with CSO-0644/0593 and sequencing with CSO-0759/3270. The fusion of *rdxA*::P*_metK_*-*cj0892c*-sfGFP was amplified from pSSv114.1 with CSO-2276/2277 and introduced into the *rdxA* locus of Δ*cj0892c* (CSS-4081) by electroporation. Clones were verified via colony PCR and sequencing with CSO-0349 and CSO-0644. Colony PCR was also used to confirm retention of the original deletion with CSO-3343 and HPK2. Similar to construction of deletion mutants, C-terminal epitope tagged strains were generated by homologous recombination at the native locus by electroporation of a DNA fragment. The 3xFLAG sequence was fused to the penultimate codon of the coding sequence to allow in-frame translation of the tag. The DNA fragment contained ∼ 500 bp upstream of the penultimate codon of the gene of interest, the sequence of the epitope tag, a non-polar resistance cassette, and the ∼500 bp downstream sequence of the gene. As an example, 3xFLAG tagging of PflA (CSS-5714) is described. The upstream fragment was amplified with CSO-4224 and CSO-4225 from *C. jejuni* NCTC11168 WT gDNA. The downstream fragment was amplified using CSO-4226 and CSO-4227. The fusion of the 3xFLAG tag with the gentamicin resistance cassette was amplified from *fliW*::3xFLAG-*aac*(3)-IV^58^ using CSO-0065 and HPK2. Next, a three-fragment overlap PCR using CSO-4224 and CSO-4227 was performed and the resulting PCR product was electroporated into CSS-4666. The obtained clones were validated by PCR using CSO-4223 and HPK2 and by sequencing using CSO-4223. PflB-3xFLAG (CSS-5716) and PflC-3xFLAG (CSS-4720) were generated similarly. The 3xFLAG with a non-polar kanamycin resistance cassette was amplified from *csrA*::3xFLAG-*aphA*-3^58^.

To construct a FlgQ-mcherry fusion protein for expression, a 76-bp DNA fragment containing the cat promoter and start codon with an in-frame BamHI site from pRY109 was amplified by PCR and cloned into the XbaI and XmaI sites of pRY112 to create pDAR1003. PCR was then used to amplify mcherry from codon to the stop codon, which was then inserted into XmaI and EcoRV sites of pDAR1003 to create pDAR1006. This plasmid contains a start codon that is in-frame with DNA for BamHI and XmaI sites followed by the mcherry coding sequence. Primers were then designed and used for PCR to amplify flgQ from codon 2 to the stop codon from C. jejuni 81-176. This fragment was inserted in-frame into the BamHI site of pDARH1006 to create pDRH7476, which was then conjugated into DRH2071 to result in DRH7516.

### CryoEM sample preparation

*C. jejuni* Δ*flhG* Δ*flaAB* cells were grown on MH plates and resuspended in phosphate-buffered saline (PBS buffer, 137 mM NaCl, 2.7 mM KCl, 10 mM Na2HPO4, 1.8 mM KH2PO4, pH 7.4). Cells were spun at 4,000 rpm for 20 min to pellet whole cells. The minicell-enriched supernatant was removed and spun in a tabletop microcentrifuge at 15,000 rpm for 5 min to pellet the minicells. The pellet was then resuspended to a theoretical OD600 of ∼15. Minicells were vitrified on QUANTIFOIL® R0.6/1 or R1.2/1.3 holey carbon grids (Quantifoil Micro Tools) using a Vitrobot Mark IV (Thermo Fisher Scientific).

For cryoET, whole-cells were grown on MHT agar, re-streaked on fresh plates and grown overnight before use. Freshly grown cells were suspended into ∼1.5 mL of PBS buffer and concentrated to an approximate theoretical OD_600_ of 10 by pelleting at 3000 rpm for 5 min on a tabletop microcentrifuge and resuspending appropriately. 30 μL of the concentrated cell sample was mixed with 10 nm gold fiducial beads coated with bovine serum albumin (BSA). 3 μL of this mixture was applied to freshly glow-discharged QUANTIFOIL® R2/2, 300 mesh grids. Grids were plunge-frozen in liquified ethane-propane using a Vitrobot mark IV.

### Single particle analysis image acquisition

Micrographs of the minicell sample were collected using 300 keV Thermo Fisher Scientific Titan Krios TEMs, across two sessions using EPU acquisition software. The first dataset was collected on a microscope with a Falcon III direct electron detector (Thermo Fisher Scientific), the second dataset using a K2 direct electron detector equipped with a GIF energy filter (Gatan), using a slit width of 20 eV. Due to our large particle size relative to that of the holes, we collected one shot per hole. Gain correction was performed on-the-fly. Details of data collection parameters are described in Table 1.

**Table 1:** Data collection statistics of two cryoEM data collection sessions.

| Parameter | Dataset 1 ("F3") | Dataset 2 ("K2") |
| --- | --- | --- |
| Voltage (kV) | 300 | 300 |
| Detector | Falcon III | K2 |
| Pixel size (Å/px) | 1.75 | 2.2 |
| Exposure dose per frame (e<sup>-</sup>/Å/frame) | 5 | 1.53 |
| Total exposure dose (e<sup>-</sup>/Å) | 50 | 50 |
| Defocus range (µm) | -1.5 to -3.0 | -1.5 to -3.0 |
| Total micrographs | 8,774 | 42,988 |

### Tilt series acquisition

Tilt series of motors in *pflA_Δ18-168_* were collected using a 300 keV Titan Krios TEM (Thermo Fisher Scientific) equipped with a K2 direct electron detector and a GIF energy filter (Gatan) using a slit width of 20 eV. Data was collected in Tomography 5 (Thermo Fisher Scientific) using a dose-symmetric tilt scheme across ±57° in 3° increments. We used a dose of 3 e-/Å^2^ per tilt, distributed across 4 movie frames. The pixel size was 2.2 Å and defocus range from - 4.0 to -5.0 μm. To determine the *C. jejuni* C-ring architecture, tilt series of 194 motors in DRH8754 were collected using a 200 keV Glacios TEM (Thermo Fisher Scientific) equipped with a Falcon 4 direct electron detector and a Selectris energy filter (Thermo Fisher Scientific) using a slit width of 10 eV. Data was collected in Leginon automated data-collection software using a dose-symmetric tilt scheme across ±51° in 3° increments. We used a dose of 2.7 e-/Å^2^ per tilt, distributed across 5 movie frames. The pixel size was 1.9 Å and defocus was -4.0 μm. All other tilt series datasets were acquired on a 200 keV FEI Tecnai TF20 FEG transmission electron microscope (FEI Company) equipped with a Falcon II direct electron detector camera (FEI Company) using Gatan 914 or 626 cryo-holders. Tilt series were recorded from −57° to +57° with an increment of 3° collected defocus of approximately −4 μm using Leginon automated data-collection software at a nominal magnification of 25,000× and were binned two times. Cumulative doses of ∼120 e−/Å2 were used. Overnight data collection was facilitated by the addition of a 3-L cold-trap Dewar flask and automated refilling of the Dewar cryo-holder triggered by a custom-written Leginon node interfaced with a computer controlled liquid nitrogen pump (Norhof LN2 Systems).

### Single particle analysis

Movie frames were aligned and dose-weighted according to exposure, as implemented in MotionCor2 version 2.1^62^. All subsequent processing was done in RELION 3.1^63,64^. CTF-correction was performed using CTFFIND4^65^, using the RELION wrapper. Flagellar motor positions were picked manually, yielding 79,287 particle coordinates for the K2 dataset and 14,605 particle coordinates in the F3 dataset.

The two datasets were first processed separately in RELION 3.1, before merging for a final round of refinement For the K2 dataset, 79287 particles were extracted at a box size of 800 px. A round of 2D classification removed junk and membrane particles, and an initial model was created using these particles with imposed C17 symmetry, which is known from past structural characterisation of the motor by subtomogram averaging^16^. A round of mask-free 3D classification and refinement with applied C17 symmetry produced the first consensus refinement. 27,164 particles were then re-extracted, centering on the periplasmic structures. After another round of 3D classification, 19,736 particles were refined in C17 symmetry to produce a whole-motor reconstruction at 9.88 Å using gold-standard refinement. For the F3 data, 14,605 particles were extracted at a box size of 1000 px rescaled to 500 px. They underwent 2D classification to remove junk, 3D classification and refinement to arrive at an initial consensus 3D structure. The particles were again re-centered on the periplasmic structures and underwent another round of refinement. Finally, the 13,054 particles were re-extracted at an un-binned 1000 px box size for a final round of refinement. The two re-centered refined datasets were merged, assigning them different RELION 3.1 optics groups, and refined to a global resolution of 9.36 Å (32,790 total particles).

Signal subtraction was used to further refine the structure of the periplasmic scaffold. A mask encompassing the regions of interest was made by segmenting and smoothing the whole-motor map using UCSF Chimera 1.16^66,67^ and its Segger plugin^68^, binarising and adding a soft-edge in RELION 3.1. The mask included the periplasmic scaffold and first ring of the basal disk, as the scaffold appears to attach onto it. This mask was used to computationally remove signal outside of the periplasmic regions of interest, as implemented in RELION 3.1. The signal subtraction and subsequent masked classification and refinement was conducted for the combined dataset, as well as K2 and F3 datasets separately. The highest resolution was reached with the merged data, the periplasmic scaffold map reaching 7.68 Å from 32,790 particles.

The periplasmic scaffold map was post-processed using LAFTER^69^ as implemented in the CCP-EM 1.6.0 software suite^70^ to suppress noise and enhance signal between the half-maps. The LAFTER-filtered map of the scaffold was used for docking and modelling of periplasmic regions.

### Focused refinement of LP-, MS- and C-rings

After convergence of the full-motor refinement in C17 symmetry, focused refinement of the MS-LP- and C-rings was carried out, using a lathed map lacking azimuthal features. C17 map was lathed by applying C360 symmetry using EMAN2, producing a map with axial and radial features but no azimuthal features. This map was used as a reference for asymmetric refinement, so that no bias towards a particular cyclic symmetry could be imposed.

For the LP-ring, a tubular mask covering the LP-ring was used for signal subtraction and focused refinement. Subtracted particles were recentred on the LP-ring in a smaller 256-pixel box. A recentred subvolume of the full lathed map was used for the initial reference. Refinement with imposition of C26 symmetry produced the LP-ring localised reconstruction.

For the MS-ring, a tubular mask covering the MS-ring was used for signal subtraction and focused refinement. Subtracted particles were recentred on the MS-ring in a smaller 256-pixel box. A recentred subvolume of the full lathed map was used for the initial reference. Refinement with imposition of C38 symmetry produced the MS-ring localised reconstruction.

For the C-ring, a toroidal mask covering the C-ring was used for signal subtraction and focused refinement against the full-size lathed motor map. Refinement with imposition of C38 symmetry produced the C-ring localised reconstruction. This map was then postprocessed and sharpened with an empirically determined B factor of -1200 Å^2^.

### Focused classification of stator complexes

From the full-motor refinement in C17 symmetry, particles were symmetry expanded and re-windowed into a smaller 360-pixel box centred on a stator complex, with subtraction of the surrounding signal using a spherical mask encompassing a triplet of neighbouring stator complexes. Classification without alignment produced the stator maps shown.

### Subtomogram averaging

Fiducial models were generated and tilt series were aligned for tomogram reconstruction using the IMOD package^71^. Tomo3D^72^ was used to reconstruct tomograms with the SIRT method due to approximately 1 particle per tomogram. All steps were automated by in-house custom scripts.

Subtomogram averaging was performed using the Dynamo package^73^ unless otherwise stated. Motors were picked using 3dmod from the IMOD suite and imported into Dynamo as ‘oriented particles’ using an in-house script, and subtomograms were extracted for averaging. For each structure, an initial model was obtained by reference-free averaging of the oriented particles, with randomized Z-axis rotation to alleviate missing wedge artefacts. This initial model was used for a first round of alignment and averaging steps, implementing an angular search and translational shifts, with cone diameter and shift limits becoming more stringent across iterations. The resulting average was used as a starting model for a round of masked alignment and averaging. In this round, custom alignment masks were implemented, focusing on the periplasmic and inner membrane-associated parts of the motor. This excluded dominant features that would otherwise drive the alignment, most prominently the outer membrane and extracellular hook. 17-fold rotational averaging was applied. The final *pflA*_Δ18-168_ average was derived from 103 particles. The Δ*pflC* average from 101 particles, Δ*pflD* average from 195 particles, and the *flgQ-mCherry* average from 155 particles.

The *C. jejuni* C-ring subtomogram average using DRH8754 used the PEET package^74^. Tomograms were CTF corrected using IMOD, motors picked using 3dmod, and subtomograms extracted for averaging in PEET. After an initial C1 whole-motor alignment, a subsequent alignment recentred on the tightly-masked C-ring was performed. To capitalise on the redundancy of C-ring archecture and to increase the number of particles, each motor was then symmetry expanded so that the C-ring beneath each of the 17 asymmetric units could be treated as a separate particle. Subsequent alignment and classification to remove poorly-aligned particles yielded the final structure.

### De novo Modelling

ColabFold^75^, the community-run implementation of Deepmind’s AlphaFold2^26^ was used to create structural models of PflA, PflB, a dimer of PflA-PflB, PflC, PflD, FlgP (as a monomer, trimer, and heptamer), FlgQ, FliL, and a dimer of MotB_C_ (with transmembrane residues 1-67 removed).

The PflAB dimer model was created by merging two predicted structures - that of a dimer of PflA residues 16-455 and PflB residues 113-820, and full-length PflA.

### Docking and refinement

For regions where α-helices and β-sheets were resolved, we refined our AlphaFold predicted protein models into the scaffold map using ISOLDE^76^ in UCSF Chimera X 1.6. We imposed torsion, secondary structure, and reference distance restraints in our modelling due to resolution limitations. The PflB chain of the PflAB dimer was rigidly docked into the map, the PflA chain was refined into the appropriate density, and any clashes resolved. Models of FlgP multimers showed that each subunit interacts with subunits i−1 and i−2. To avoid artefacts due to this, a heptamer of FlgP was docked into the innermost basal disk ring and then refined into the map. Two subunits were then removed from each end, resulting in a fitted FlgP trimer. The PflC model was first separated into three domains, PflC_N_ (residues 16-252), linker (253-263), and PflC_C_ (264-364). Each domain was rigidly docked into its appropriate density, the three chains merged, and the resulting protein was fitted using ISOLDE to resolve clashes or poor geometry at merging points. Six copies of PflC_N_ were rigidly docked into an asymmetric unit of PflC lattice in the medial disk, forming PflC_2-7_. PflD, MotB, and an arc of FliL, were rigidly docked into the LAFTER-filtered scaffold map.

We used *phenix.validation_cryoem* from the Phenix package^77^ to evaluate map and model quality for PflA, PflB, PflC, PflC_2-7_, PflD, FlgP trimer, MotB dimer, and FliL. For each model, a soft mask of the region was first created and applied to the map. The resulting masked volume and corresponding structural model were input to *phenix.validation_cryoem* which generated map-model FSC plots or cross-correlation scores.

### Flagellar motility assays

WT *C. jejuni* and DAR2076 were grown from freezer stocks on MH agar containing trimethoprim for 48 h in microaerobic conditions at 37 C. Strains were restreaked on MH agar containing trimethoprim and grown for 16 h at 37 C in microaerobic conditions. After growth, strains were resuspended from plates and diluted to an OD_600_ 0.8. Strains were then stabbed in MH motility agar (0.4% agar) and incubated at 37 C in microaerobic conditions for 30 h and assessed for migration from the point of inoculation.

### Bead assay of rotational steps

The *C. jejuni* bead assay strain was constructed by deleting *flaA* to shorten the flagellar filament, making a cysteine substitution in *flaB* to allow for filament bi-otinylation via a malemide-PEG2-biotin linker (Thermo Scientific A39261), making an alanine substitution in *flhF* which moves motors to a subpolar position permitting better bead coupling, and deleting *cheY* to prevent motor switching. From frozen stocks, this strain was cultured on Mueller-Hinton agar (1.4% w/v) supplemented with trimethoprim (10 μg/mL) overnight at 37 ◦C in a pouch with a gas-controlling sachet (EZ CampyPouch System BD 260685) and a damp paper towel. The growth from this culture was restreaked onto another plate and grown in the same conditions overnight. Passages to new plates on the following two days were optionally made to propagate the culture for multiple days of experiments.

After growth, bacteria were washed off the plate with MH broth and washed twice with PBS (centrifuged for 2 min at 11 500 RPM) to create a 500 μL suspension of OD_600_ of 2. Next, 25 μL of 20 mM malemide-PEG2-biotin linker in PBS was added and left to react for 10 min. The bacteria were then washed twice more in PBS. A custom tunnel slide composed of a slide with two drilled holes, a parafilm spacer, and another glass slide was constructed and connected to a peristaltic pump (Lambda Multiflow) to draw flow through the slide. Poly-L-lysine (PLL) solution (Sigma-Aldrich P4707) was flowed into the slide channel and left for 5 min to coat the glass surface. Excess PLL was flushed with PBS. Bacteria were added and left 5 min to settle on the surface. Unattached bacteria were flushed sequentially with PBS and MH broth. 1300 nm diameter, streptavidin-coated beads (Sigma-Aldrich 49532) were washed once in PBS and dispersed in MH broth. Beads were added to the flow slide and left for 5 min to settle and conjugate to filaments. Excess beads were flushed away with MH-broth. Spinning beads were observed with a bright-field microscope with a high numerical aperture (NA) objective (100×, 1.46 NA, Zeiss 420792-9800-720) and recorded with a high speed scientific complementary metal-oxide semiconductor camera (Optronis CL600X2-FULL-M-FM). Positions of slowly-rotating beads attached to cells treated with carbonyl cyanide 3-chlorophenylhydrazone (CCCP) at concentrations of 0 μM to 5 μM in MH broth revealed preferred phase-invariant dwell positions.

### Plasmid construction and cloning in E. coli

*C. jejuni* proteins for recombinant expression in *E. coli* were cloned into the pLIC plasmid backbone, which confers resistance to ampicillin and places the gene of interest under an IPTG-inducible T7 promoter for high levels of controlled expression. We used WT *C. jejuni* genomic DNA as template for gene amplification (extracted using the Wizard genomic DNA purification kit by Promega), and the Gibson Assembly method^78^ to seamlessly assemble all plasmid constructs. For all constructs, primer pairs were designed to amplify 1) the pLIC backbone and 2) the gene to be expressed, while also introducing a 25∼30 bp complementary overlap between the two fragments. The pLIC plasmid primers also introduced an N-terminal hexahistidine tag. After vector linearisation and purification of PCR product, it was digested with DpnI (New England Biolabs) to remove template vector.

The resulting linear DNA fragments were assembled using the Gibson Assembly master mix (New England Biolabs). 5 μL of mix was added to 15-20 fmol of linearised vector and 4× excess of insert and topped up to 10 μL with double-distilled water (ddH_2_O). The tube was incubated at 50 C for 15 min and kept on ice until transformation.

Before transformation, 30 μL of ddH_2_O was added to the 10 μL reaction. 2 μL of diluted Gibson mix was added to 25 μL of chemically competent *E. coli* DH5α and transformed using the heat shock method^79^. The entire volume of the tube was then plated onto a carbenicillin-supplemented LB agar plate.

After confirmation by Sanger sequencing (Source Bioscience), each assembled construct was isolated from the cloning strain (QIAprep Spin Miniprep Kit, QIAGEN) and transformed into E. coli BL21(DE3) for recombinant overexpression.

### Protein overexpression and purification

All proteins encoded on pLIC expression vectors were purified using the same protocol. A small (5 mL) overnight liquid culture of *E. coli* BL21(DE3) carrying the appropriate expression vector was prepared and diluted 1:50 in 1000 mL of LB medium. Shaking at 37 C, the culture was grown to OD600 0.4-0.6, after which protein expression was induced by addition of 0.5 mM IPTG. Temperature was reduced to 18 C and protein was expressed overnight.

Cells were harvested at 5000 rpm, 4 C for 20 min. All subsequent steps were done on ice using buffers chilled to 4 C. The cell pellet was gently resuspended in ∼35 mL of wash buffer (50 mM Tris-HCl, 100 mM NaCl, 30 mM imidazole, pH 7.5). DNase and protease inhibitor were added (cOmplete Protease Inhibitor Cocktail, Roche). Cells were lysed using a LM10 Microfluidizer Processor cell disrupter (Analytik) at 15,000 psi. Lysate was centrifuged at 17,000 rpm, 4 C for 30 min to pellet debris. The resulting supernatant was filtered through a 0.45 μm syringe filter (Whatman).

A 5 mL HisTrap HP affinity chromatography nickel column (Cytiva) was first equilibrated with wash buffer. Supernatant was loaded onto the column with a peristaltic pump at a flow rate of 3 mL/min. The column was washed with 50 mL of wash buffer and then transferred onto a Fast protein liquid chromatography system (BioRad). The column was further washed until the UV trace was flat. Then, protein was eluted from the column using a high-imidazole buffer (50 mM Tris-HCl, 100 mM NaCl, 500 mM imidazole, pH 7.5) at a flow of 2 mL/min using ‘reverse flow’.

The purified protein was kept at 4 C or flash-frozen in LN2 for longer-term storage before characterising them by mass photometry.

### Analytical size exclusion

Analytical size exclusion of PflC and PflC_N_ (Δ236-349) was performed with a ENrich SEC 650 column (Bio-Rad), equilibrated with 1× PBS at a flow rate of 0.1 ml min−1 and a total sample injection volume of 400 µl. The column was calibrated using the Protein Standard Mix 15 – 600 kDa (Supelco #69385). Absorption was recorded at 280, 220 and 495 nm to follow elution profiles and plotted using GraphPad Prism.

### Protein pulldowns

#### Co-immunoprecipitation (coIP) of PflA/B-3xFLAG with PflD-sfGFP, and PflC-3xFLAG

Chromosomally epitope-tagged fusions of PflC-3xFLAG (CSS-4720) or PflA-3xFLAG and PflD-sfGFP (CSS-5714), and PflB-3xFLAG and PflD-sfGFP (CSS-5716) were used together with the untagged *C. jejuni* NCTC11168 WT (CSS-0032) and PflD-sfGFP only (CSS-4666) as controls for immunoprecipitation. Co-purification of FlgP or PflD-sfGFP was investigated by western blot (WB) analysis using FlgP specific antisera^19^ or an anti-GFP antibody (Roche #11814460001, RRID:AB_390913), respectively. In brief, strains were grown to an OD_600_ of 0.6 and 60 OD_600_ of cells were harvested (5,000 rpm, 20 min, 4 C) and washed in buffer A (20 mM Tris-HCl pH 8, 1 mM MgCl_2_, 150 mM KCl, 1 mM DTT). In parallel, 1 OD_600_ of cells was harvested as “culture” control and boiled in 1 x protein loading buffer (PL; 62.5 mM Tris-HCl, pH 6.8, 100 mM DTT, 10% (v/v) glycerol, 2% (w/v) SDS, 0.01% (w/v) bromophenol blue; 8 min at 95 C, shaking at 1,000 rpm). Next, 60 OD_600_ cell pellets were lysed with a FastPrep system (MP Biomedical, matrix B, 1 x 4 m/s, 10 s) in 1 ml lysis buffer [buffer A including 1 mM PMSF (phenylmethylsulfonyl fluoride, Roche), 20 U DNase I (Thermo Fisher Scientific), 200 U RNase Inhibitor (moloX, Berlin) and Triton X-100 (2 µl/ml lysis buffer)]. Cleared lysates (13,000 rpm, 10 min, 4 C) were incubated with 35 µl anti-FLAG antibody (Sigma-Aldrich, #F1804-1MG, RRID:AB_262044) for 30 min at 4 C with rotation. Before and after incubation, a 1 OD_600_ aliquot was taken aside as lysate and supernatant 1 samples. Lysates with anti-FLAG antibody were then incubated for additional 30 min (4 C, rotating) with 75 µl/sample pre-washed (3 times in buffer A) Protein A-Sepharose beads (Sigma-Aldrich, #P6649). Afterwards, the supernatant/unbound fraction was removed after centrifugation (15,000 x g, 1 min, 4 C; supernatant 2) and Protein A-Sepharose beads with bound proteins were washed 5 times with buffer A. Elution of the bound proteins was performed with boiling of the beads in 400 µl 1 x PL (8 min at 95 C, 1,000 rpm). Six volumes of acetone were used to precipitate eluted proteins overnight at -20 C. Next, precipitated proteins were harvested by centrifugation (15,000 rpm, 1 h, 4 C), air-dried and resuspended in 1 x PL. Culture, lysate, supernatants 1 & 2, wash (aliquots corresponding to 0.1 OD_600_) and eluate samples (corresponding to 10 OD_600_) were analysed by WB. Western Blots were performed as described previously^58^ and probed with the appropriate primary antibodies (anti-FLAG, anti-GFP (1:1,000 in 3% BSA/TBS-T)) or FlgP antisera (1:20,000 in 3% BSA/TBS-T) and secondary antibodies (anti-mouse or anti-rabbit IgG, HRP-conjugate (1:10,000) in 3% BSA/TBS-T; GE Healthcare, #RPN4201 and #RPN4301, respectively).

### Mass Photometry

Microscope coverslips (24 × 60 mm, Carl Roth) and CultureWell Gaskets (CW-50R-1.0, 50-3 mm diameter × 1 mm depth) were cleaned with alternating ddH_2_O and 100% isopropanol washes, then dried roughly with pressurised air and left to dry further overnight at room temperature. Before use, gaskets were assembled onto coverslips and placed on the lens of a One^MP^ mass photometer (Refeyn Ltd) with immersion oil.

For each measurement, a gasket was first filled with 18 μL of PBS buffer and the instrument was focused. Then, 2 μL of sample was added to the droplet and rapidly mixed by pipetting. Measurements were then started using AcquireMP v1.2.1 (Refeyn Ltd). For each measurement, data was acquired for 60 s at 100 frames per second. Mass photometry data was processed and analyzed in DiscoverMP software v.1.2.3 (Refeyn Ltd).

Measurements were conducted using affinity chromatography-purified proteins diluted to 200-800 nM, calculated from absorption at 280 nm. 2 μL of sample was added to 18 μL PBS droplet and mixed. For measurements of hetero-oligomers, the different proteins were first combined and mixed in a separate tube and subsequently applied to the PBS droplet. MP measurements were calibrated against molecular masses of commercial NativeMark^TM^ unstained protein standard (Thermo Fisher Scientific). 1 μL of NativeMark^TM^ was diluted 30-fold in PBS and 2 μL of this solution was added to 18 μL PBS for measurement. Detected peaks corresponded to 66 kDa, 146 kDa, 480 kDa, and 1048 kDa and were used to calibrate subsequent measurements in DiscoverMP.

### Construction of plasmids and strains for PflA and PlfB co-immunoprecipitation experiments

Plasmids were constructed with specific promoters for expression of FLAG-tagged proteins in *C. jejuni* mutants for co-immunoprecipitation experiments. To express a C-terminal FLAG-tagged PflA protein, a 206-base pair DNA fragment from *C. jejuni* 81-176 that contained the promoter for *flaA* encoding the major flagellin with its start codon and an in-frame SpeI restriction site followed by an in-frame BamHI restriction site was amplified by PCR. This fragment was cloned into the XbaI and BamHI sites of pRY108 to result in pDAR1425. Primers were then constructed to amplify DNA from codon 2 to the penultimate codon of *pflA* from *C. jejuni* 81-176 with an in-frame C-terminal FLAG tag epitope and stop codon. This DNA fragment was then cloned into the BamHI site of pDAR1425 so that *pflA*-FLAG was expressed from the flaA promoter to create pDAR3417. As a control, a 229-base pair DNA fragment from *C. jejuni* 81-176 that contained the promoter for *flaA* encoding the major flagellin with its start codon and DNA encoding an in-frame FLAG tag epitope followed by an in-frame BamHI restriction site was cloned into the XbaI and BamHI sites of pRY108 to create pDAR1604. pDAR1604 and pDAR3417 were then moved into DH5a/pRK212.1 for conjugation into DAR1124. Transconjugants were selected for on media with kanamycin and verified to contain the correct plasmids to result in DAR3447 and DAR3477.

To express a C-terminal FLAG-tagged PflB protein, primers were constructed to amplify DNA from codon 2 to the penultimate codon of pflB from *C. jejuni* 81-176 with an in-frame C-terminal FLAG tag epitope and stop codon. This DNA fragment was then cloned into the BamHI site of pECO102 so that pflB-FLAG was expressed from the cat promoter to create pDAR3414. pDAR965 and pDAR3414 were then moved into DH5a/pRK212.1 for conjugation into DAR981. Transconjugants were selected for on media containing chloramphenicol and verified to contain the correct plasmids to result in DAR3451 and DAR3479.

### PflA and PflB co-immunoprecipitation experiments

*C. jejuni* Δ*pflA* and Δ*pflB* mutants containing plasmids to express a FLAG-tag alone or C-terminal FLAG-tagged PflA or PflB proteins were grown from freezer stocks on MH agar containing chloramphenicol for 48 h in microaerobic conditions at 37 C. Each strain was restreaked onto two MH agar plates containing chloramphenicol and grown for 16 h at 37 C in microaerobic conditions. After growth, strains were resuspended from plates in PBS and centrifuged for 10 min at 6000 rpm. Each cell pellet was resuspended in 2 ml of PBS. Formaldehyde was added to a final concentration of 0.1% and suspensions were gently mixed for 30 min at room temperature to crosslink proteins. After crosslinking, 0.4 ml of 1 M glycine was added to each sample and then suspensions were gently mixed for 10 min at room temperature to quench the crosslinking reaction. Bacterial cells were collected by centrifugation for 10 min at 6000 rpm. Cells were then disrupted by osmotic lysis and FLAG-tagged proteins with associated interacting proteins were immunoprecipitated with α-FLAG M2 affinity resin as previously described^80,81^.

To identify potential proteins interacting with PflA and PflB, resin with immunoprecipitated proteins were resuspended in SDS-loading buffer and electrophoresed on a 4-20% TGX stain-free gel (Bio-Rad) for 10 min. The gel was then stained with Coomassie blue for 30 min and then destained overnight. After equilibration of the gel in dH_2_O for 30 min, a 1 cm region of the gel containing a majority of the co-immunoprecipitated proteins was excised and diced into 1 mm pieces and then submitted for analysis by LC-MS/MS. After identification of proteins that co-immunoprecipitated with the FLAG-tagged bait protein and with the resin from the FLAG-tag only sample (the negative control), a ratio for each protein was determined by dividing the abundance of each protein detected in the FLAG-tagged bait protein sample by the abundance of each protein in the negative control. The top twenty proteins with the highest ratios for co-immunoprecipitation with the FLAG-tagged bait proteins are reported. The top twenty proteins that only co-immunoprecipitated with the FLAG-tagged bait proteins and were not detected in the negative control samples are also reports with their respective raw abundance counts.

## Extended data figure legends

**Extended Data Figure 1.**
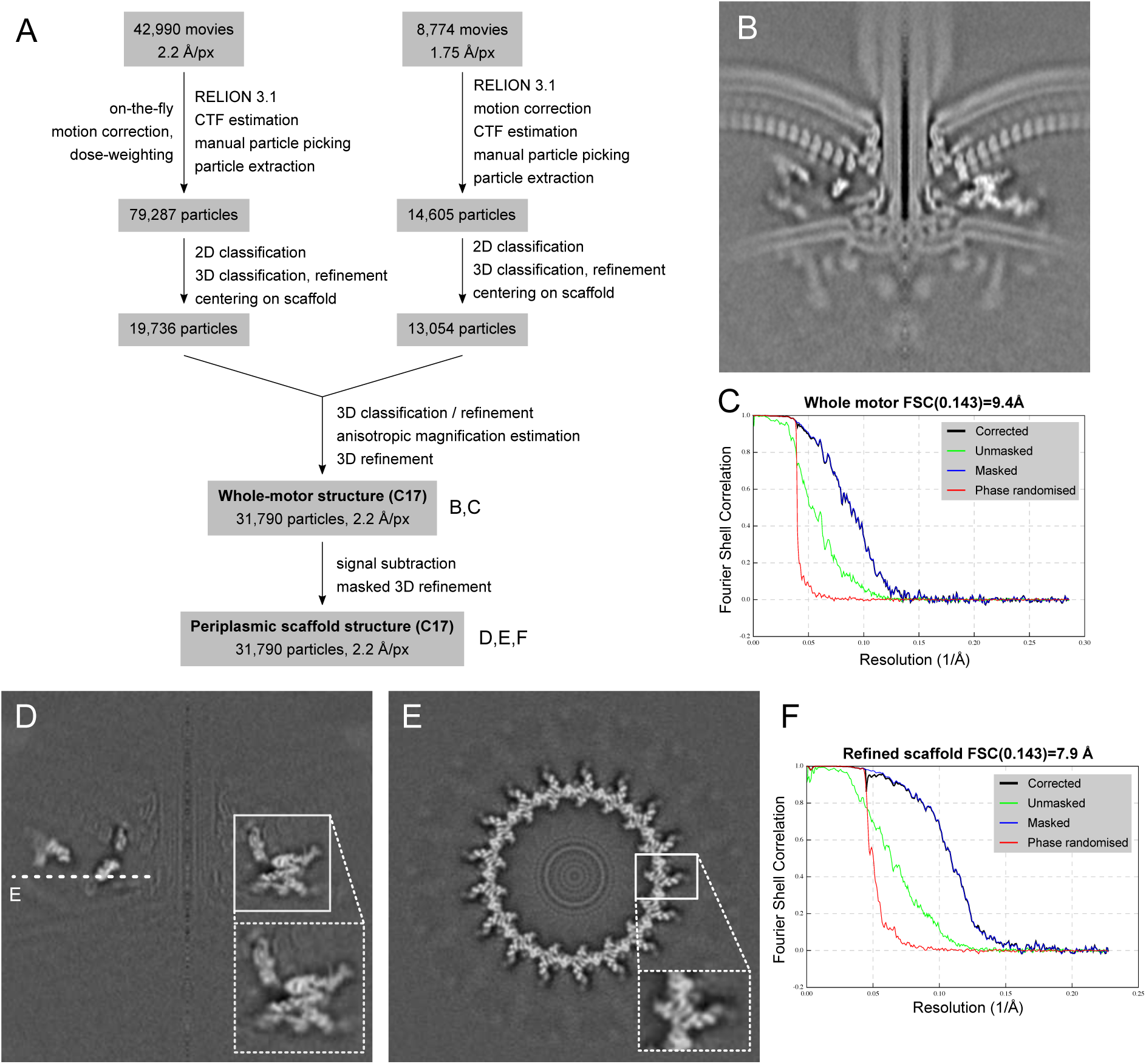
Flowchart and resolution estimates of structure determination of the *Campylobacter jejuni* bacterial flagellar motor using in situ single particle analysis. (A) Simplified flowchart showing the generation of cryoEM volumes. (B) Central slice through the refined whole-motor structure. (C) FSC curve for B. (D) and (E) show slices through the volume of the refined, signal-subtracted periplasmic scaffold. (F) FSC curve for D, E.

**Extended Data Figure 2.**
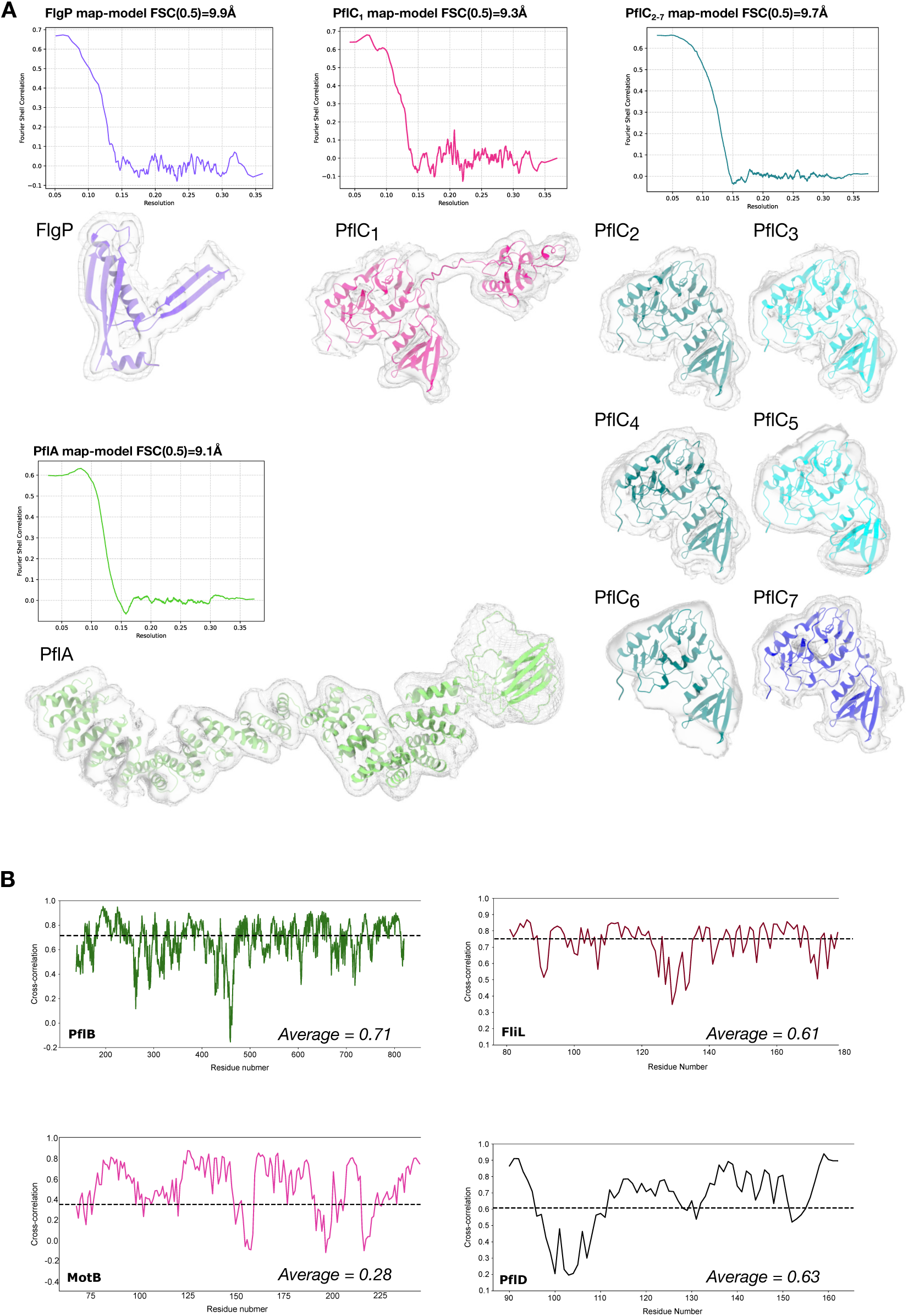
Validation of those protein chains modelled in the scaffold map to subnanometre resolution. (A) Map-model FSC curves for protein models refined into the scaffold map: FlgP, PflA, PflC_1_, and PflC_2-7_ calculated in Phenix, alongside images of excised pieces of the density map corresponding to individual proteins. High and low isosurface thresholds are denoted by solid and mesh surfaces, respectively, to illustrate fit of models into secondary structure densities. (B) Cross-correlation per residue plots of proteins docked into the map: PflB, FliL, MotB, PflD, calculated using Phenix. Mean CC values are shown with dashed line and in text inset.

**Extended Data Figure 3.**
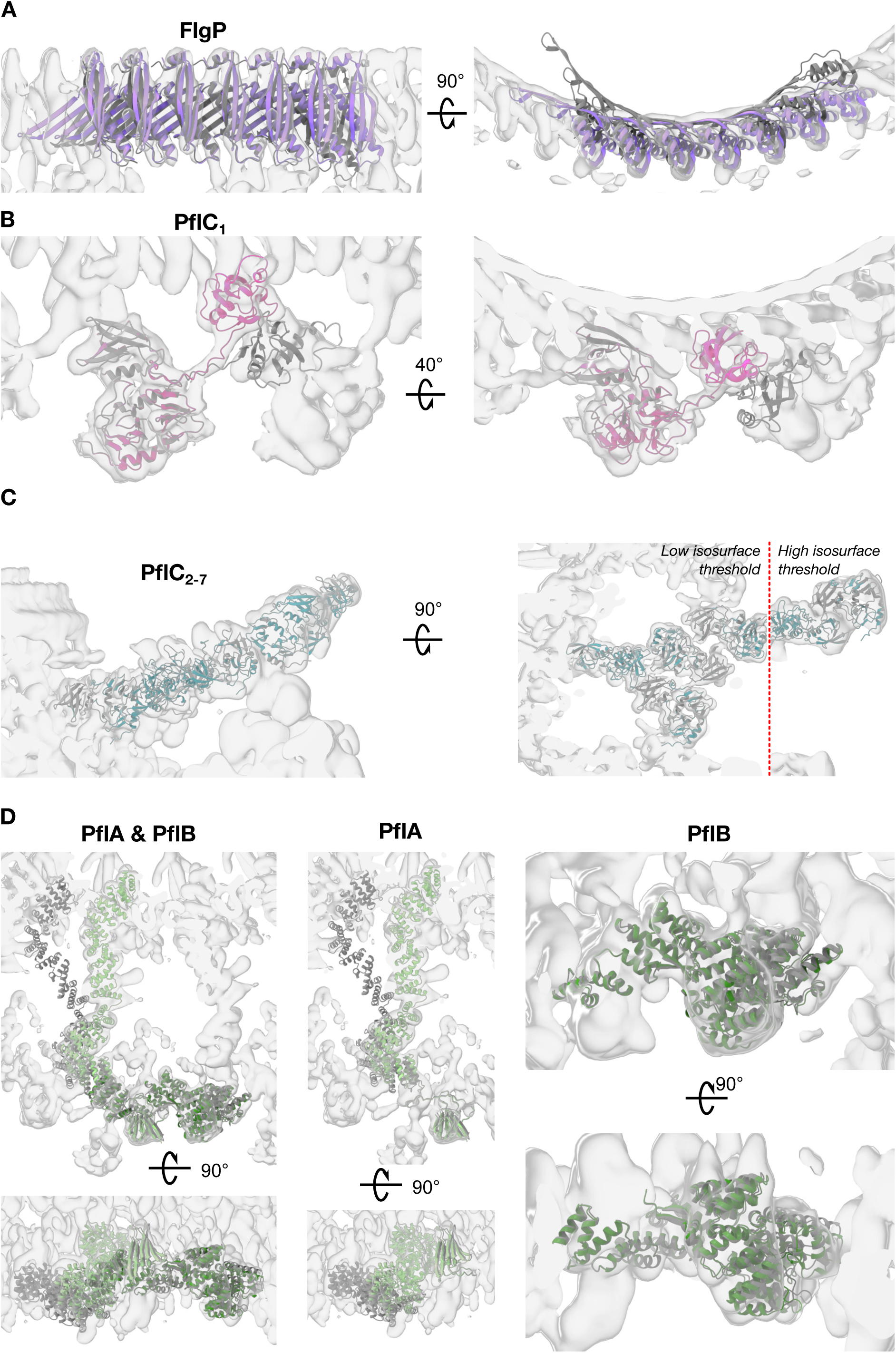
Comparison of AlphaFold2 starting models with final structures after flexible fitting. (A) AlphaFold2 prediction of FlgP (dark grey) required bending of the oligomer to fit the curvature of the first ring of the basal disk (purple). (B) AlphaFold2 prediction of PflC_1_ required independent rigid-body docking of the N- and C-terminal domains (magenta). (C) The resulting N-terminal domain of PflC was rigid-body docked into density multiple times for PflC_2-7_. C-terminal domains were not modelled. Higher thresholds required for more peripheral, and presumably more flexible, components. (D) PflA and PflB AlphaFold2 models (dark grey) required bending to fit into density maps (pale and dark green for PflA and PflB, respectively).

**Extended Data Figure 4.**
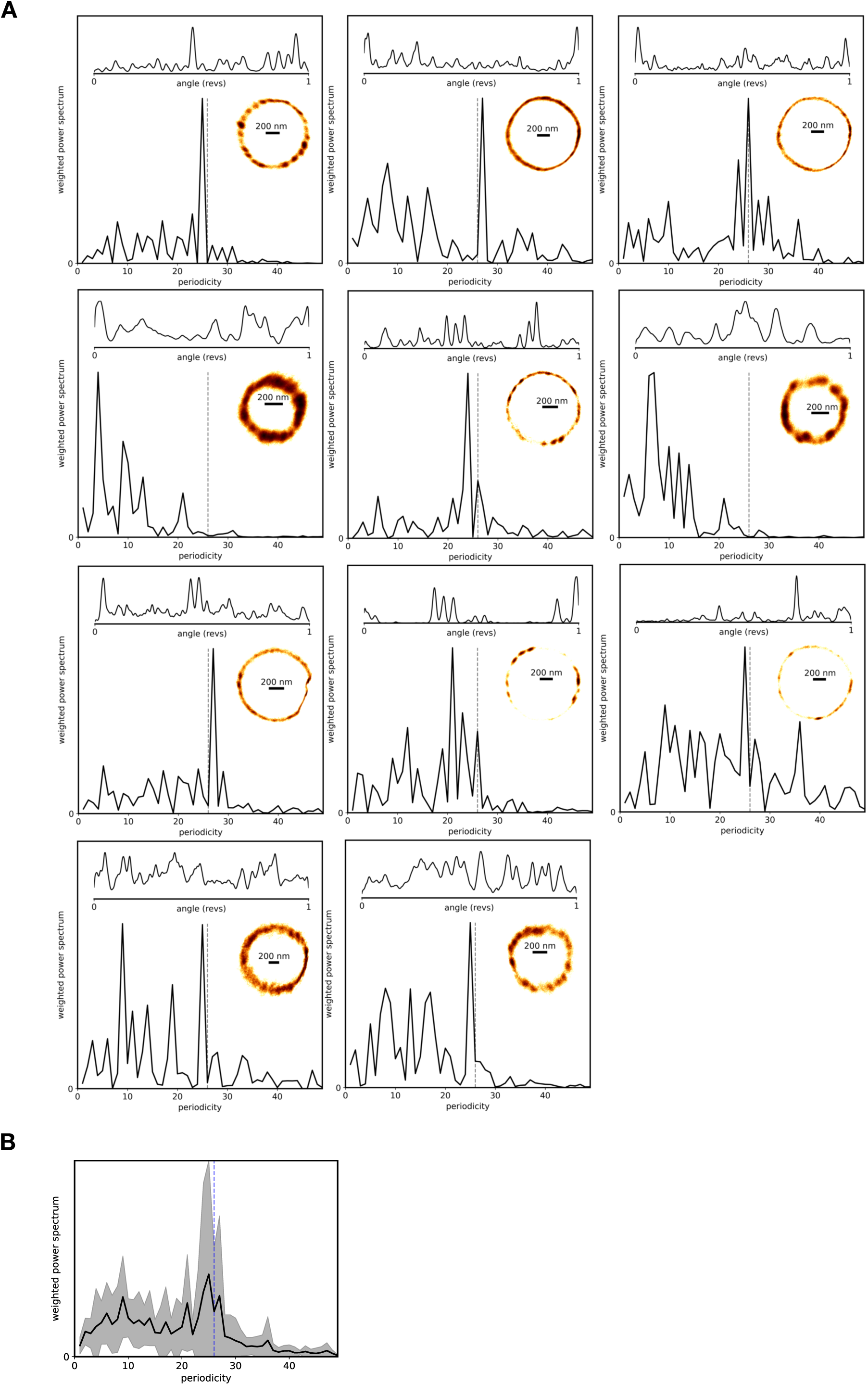
Discrete steps in flagellar rotation are similar to those in *E. coli.* (A) Kernel density estimation bead position plots over many rotations for 11 cells. Measurements were of duration 20 to 240 seconds, substantially de-energized by CCCP to slow rotation. Their (x,y) histograms, a kernel density of the angular position, and the weighted power spectrum are depicted. The top left is the trace shown in Fig. 2. (B) Weighted power spectrum of all 11 traces.

**Extended Data Figure 5.**
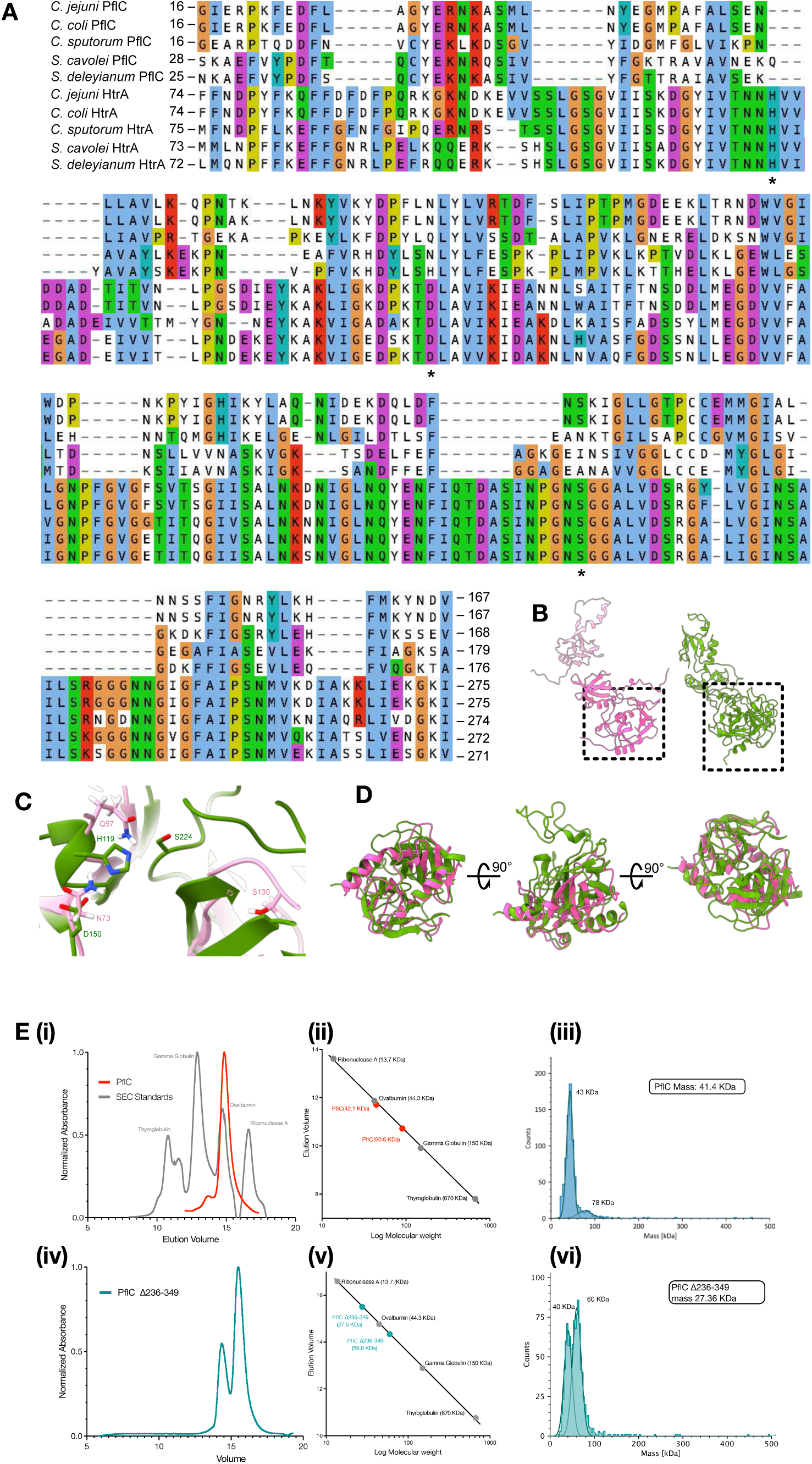
PflC (a previously unknown glycoprotein) resembles an inactivated HtrA that forms oligomerises *in vitro,* regulated by its C-terminal PDZ domain. (A) Multiple sequence alignment of PflC and HtrA protease domains from species in the sister clades *Campylobacter* and *Sulfurospirillum*. Alignment is based on structural alignments of PflC and HtrA models, with local sequence re-alignments. Asterisks highlight catalytic triad residue locations that are conserved as His-Asp-Ser in HtrA but not PflC. (B) Structurally aligned PflC (left) and HtrA (right), highlighting location of the protease domain. (C) The His-Asp-Ser catalytic triad in the protease domain of HtrA (green) is not conserved in PflC (pink). (HtrA PDBID: 6Z05). (D) Structural alignment of the HtrA domains of HtrA and PflC in ChimeraX Matchmaker. RMSD between pruned atom pairs 1.3 Å, across all atoms 10.4 Å. (E) Oligomerisation of PflC is regulated by its C-terminal domain: (i) Size Exclusion Chromatography of PflC (red) along with protein standards (grey) showing elution volume. (ii) Calibration graph showing a dimer of PflC (red) corresponding to retention times during elution. (iii) Mass photometry measurements of purified PflC (replicates). (iv) Size Exclusion Chromatography of PflCN (Δ236-349, green) along with protein standards (grey) showing elution volume. (v) Calibration graph of PflCN (green) corresponding to retention times during elution. (vi) Mass photometry measurements of purified PflCN. Theoretical masses are shown in the inserts. The instrument’s limit of detection is 35 kDA, meaning the monomer mass is larger than otherwise expected.

**Extended Data Figure 6.**
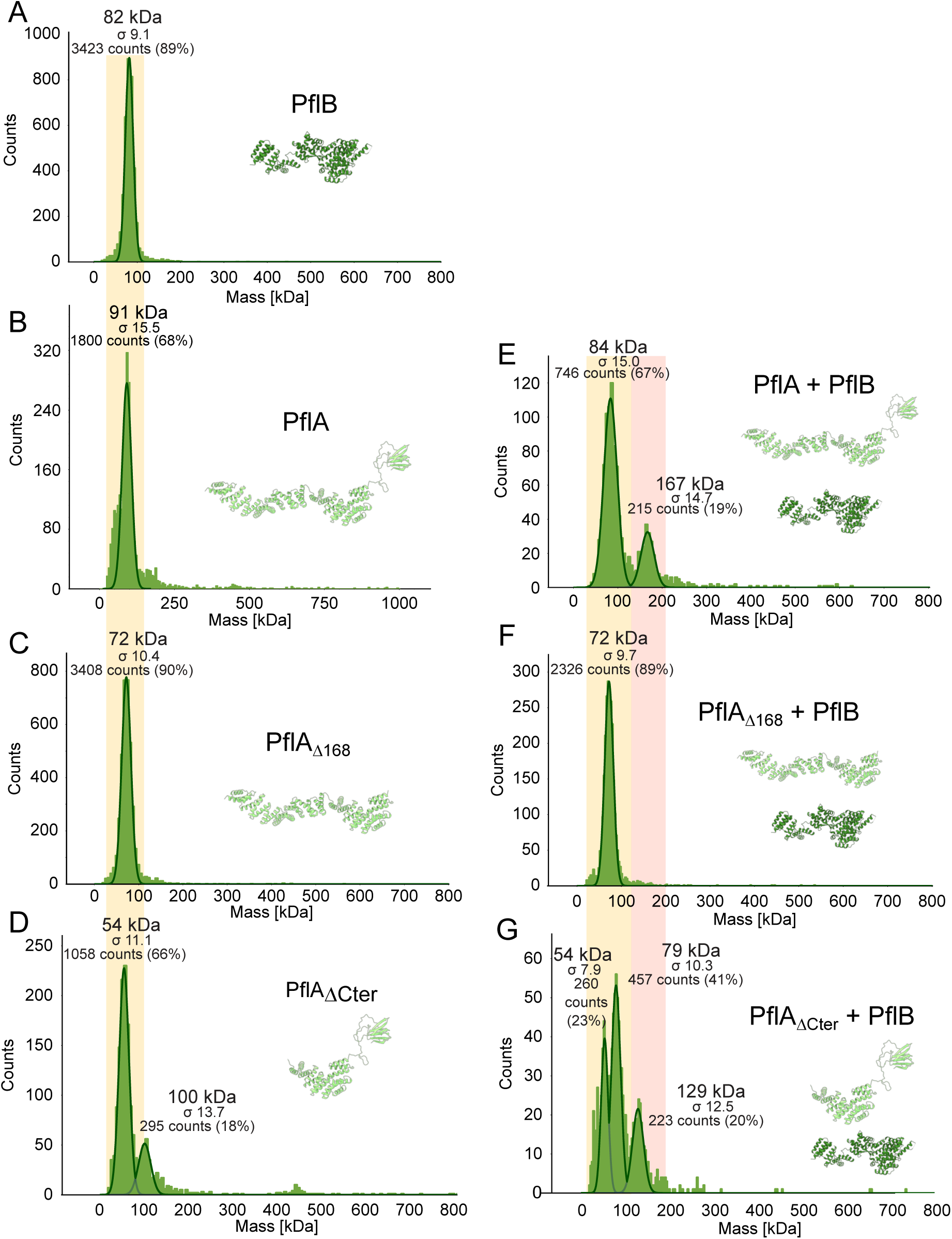
Mass photometry shows that PflA dimerises with PflB via its N-terminal β-sandwich domain. (A-D) Mass photometry measurements of purified PflA and PflB constructs show the proteins are mainly monodisperse. There is a dimer peak present for the PflA_ΔCter_ construct, likely due to a reduction in stability and solubility. (E-G) Mass photometry measurements of mixtures of PflB and PflA variants. Dimer peaks appear only when β-sandwich and linker domain of PflA is present. In panels E and G, monomer peaks of PflA and PflB are not resolved due to their similar molecular weights. In the bottom panel, the 54 kDa peak corresponds to PflA_ΔCter_, and the 79 kDa peak to PflB. Broadly, monomer peaks have a yellow background, dimer peaks red.

## Supplemental table titles

**Table S1:**
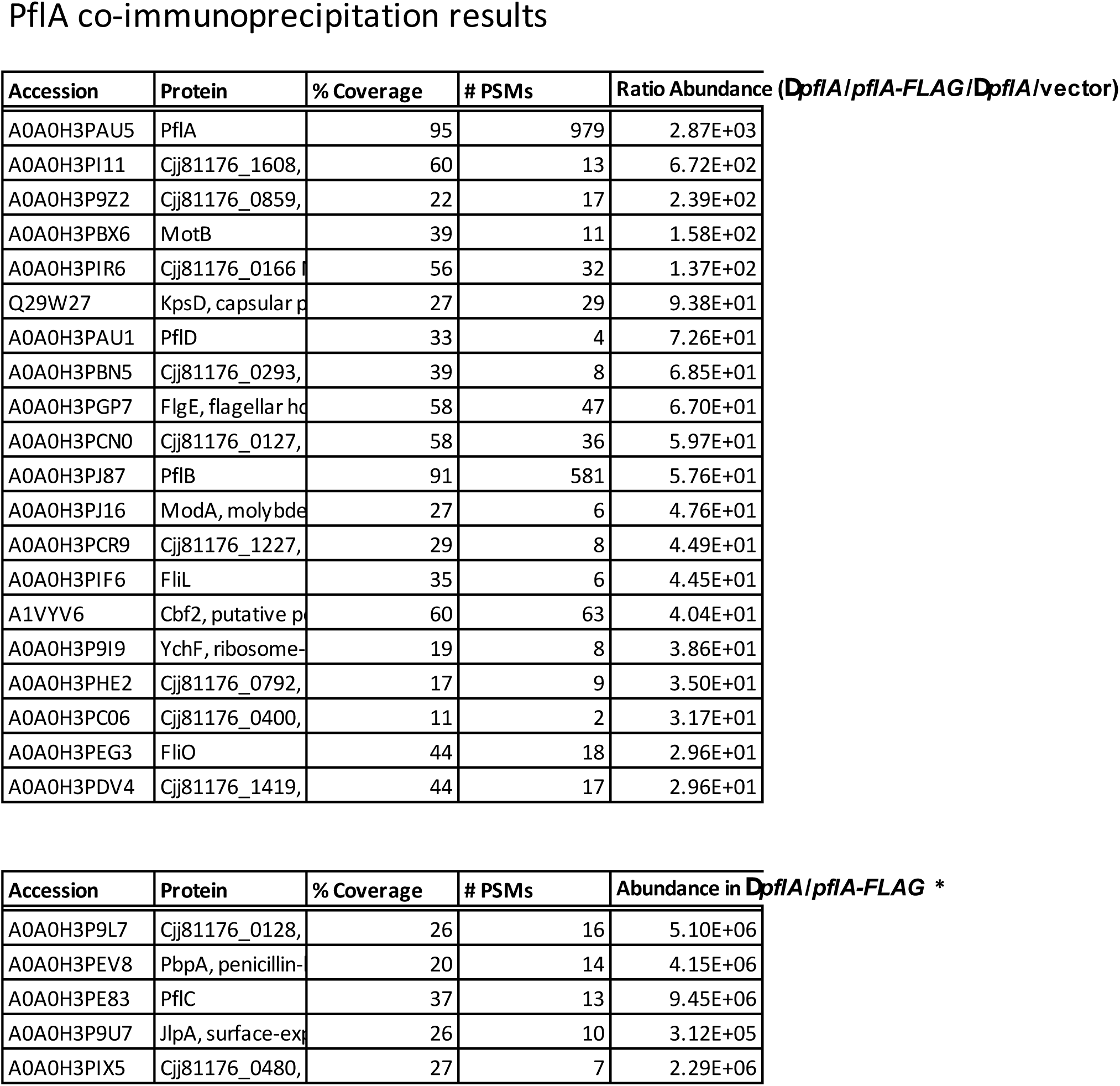
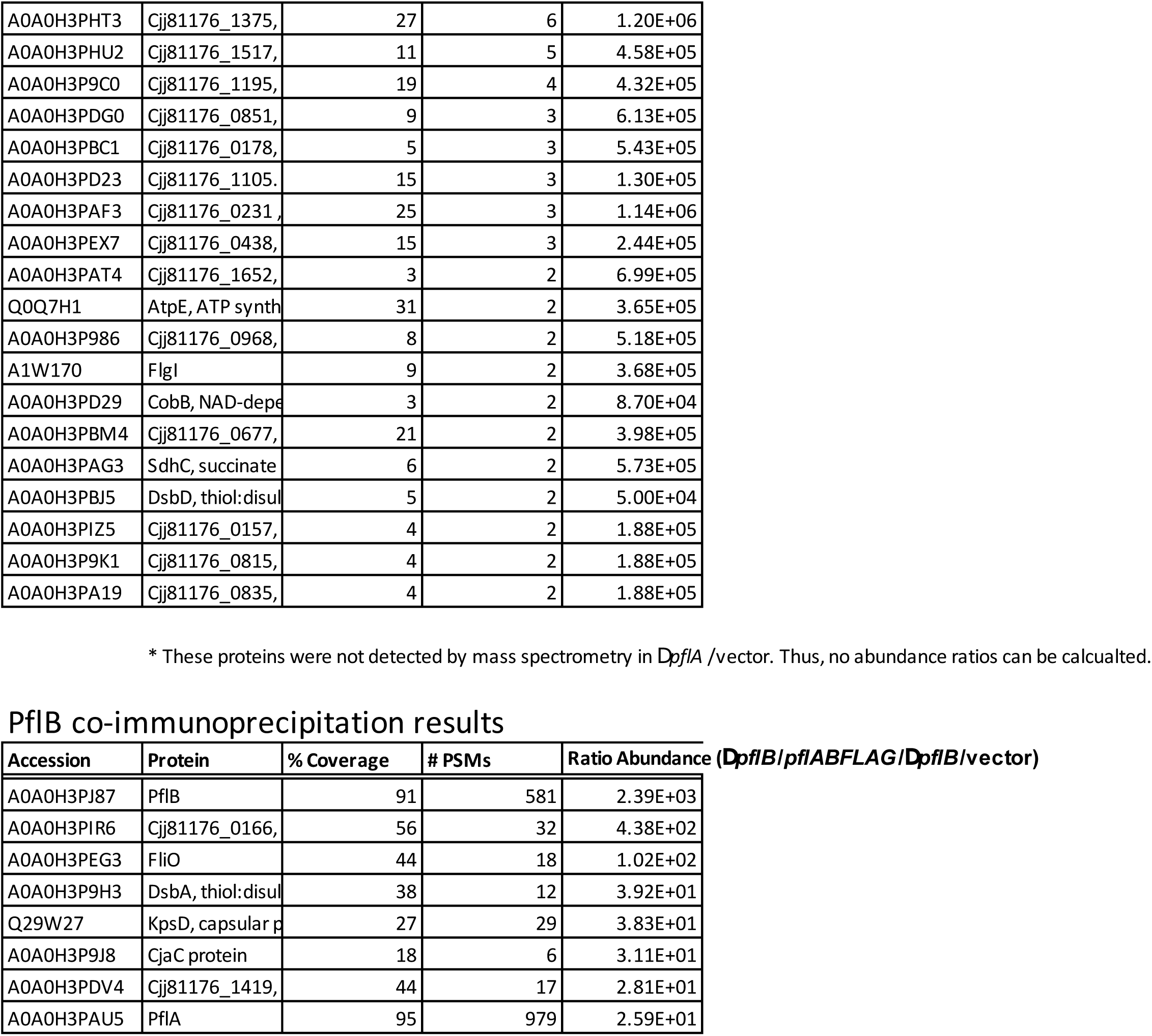
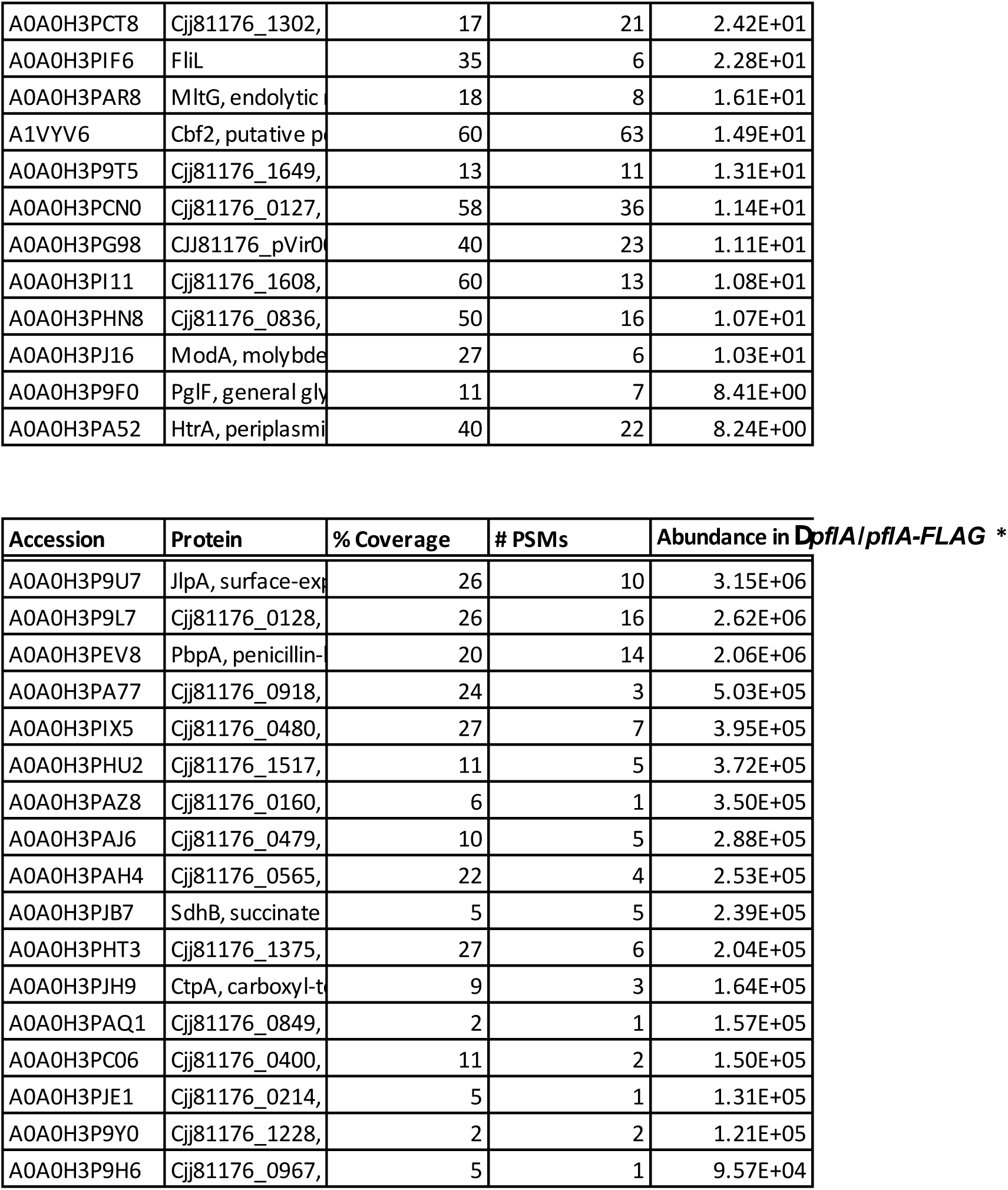
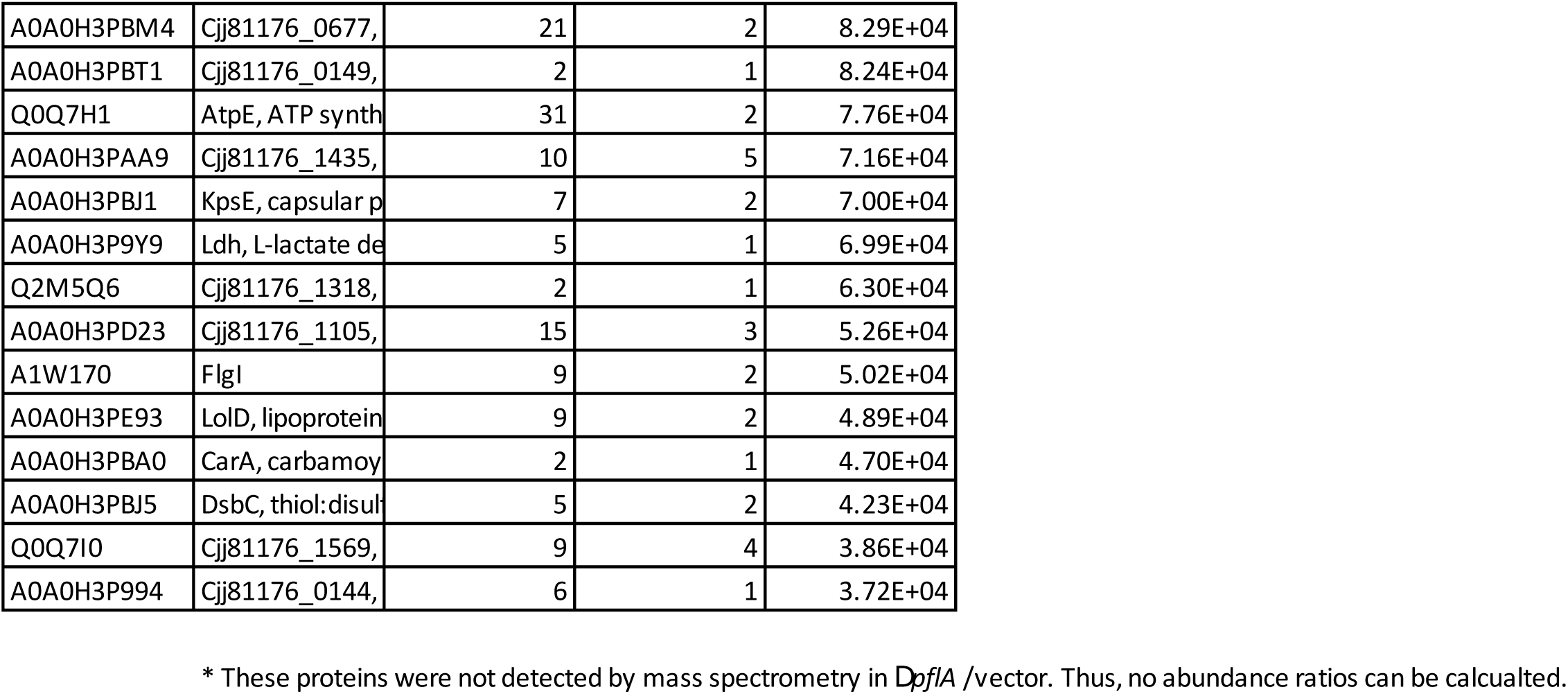
PflA and PflB pulldowns and mass spectrometry.

**Table S2:** Structural components of the *Campylobacter jejuni* flagellar motor.

| **Protein** | **Accession number** | **Functional description** | **Color in figures** | Approx. resolution (d99) cite PMID 30198894 | **Stoichiometry** | **Region modelled** | **Structural source** |
| --- | --- | --- | --- | --- | --- | --- | --- |
| MotB | A0A0H3P BX6 | Stator unit component | Light pink | >10 | 2 (x17) | 15-55 (TM) 68-247 (periplasmic) | From PMID 32931735 AlphaFold2 |
| FliL | A0A0H3PI F6 | MotB-associated periplasmic protein | Dark red | >10 | 4 (x17) | 81-178 | Homology model from PMID 35046042 |
| PfIA | A0A0H3P AU5 | Scaffold protein that recruits PfIB | Light green | 9.1 | 1 (x17) | 16-788 | AlphaFold2 |
| PfIB | A0A0H3P J87 | Scaffold protein that recruits stator complexes by MotB | Dark green | >10 | 1 (x17) | 138-820 | AlphaFold2 |
| PfIC | A0A0D7VI M6 | Scaffold protein that attaches the scaffold to the basal disk, makes up the medial disk | Magenta, teal | 9.3 - 9.7 | 1 (x17)+<br>6 (x17) | 16-364 | AlphaFold2 |
| PfID | A0A0H3P AU1 | Scaffold protein | Black | >10 | 1 (x17) | 91-162 | AlphaFold2 |
| FlgP | A0A0H3P CP8 | Makes up the basal disk, a large rigid OM-associated brace | Purple | 9.9 | 51 | 66-171 | AlphaFold2 |

**Table S3:** Antisera, strains, and recombinant DNA used in this study.

| REAGENT or RESOURCE | SOURCE | IDENTIFIER |
| --- | --- | --- |
| Antibodies |  |  |
| FlgP specific antisera | Ref<sup>21</sup> | FlgP specific antisera |
| Anti-GFP | Roche | #11814460001, RRID:AB_390913 |
| Anti-FLAG | Sigma-Aldrich | #F1804-1MG, RRID:AB_262044 |
| Anti-mouse IgG | GE Healthcare | #RPN4201 |
| Anti-rabbit IgG | GE Healthcare | #RPN4301 |
| Bacterial and virus strains |  |  |
| *C. jejuni* 81-176 *rpsL*<sup>Sm</sup> (Sm<sup>R</sup>) | Ref<sup>53</sup> | *C. jejuni* DRH212 |
| *C. jejuni* 81-176 $\Delta flhG \Delta flaAB$ | This work | *C. jejuni* minicell-producing strain |
| *C. jejuni* 81-176 *pflA*<sub>$\Delta 18-168$</sub> | This work | *C. jejuni* PflA truncation |
| *C. jejuni* NCTC11168 wildtype | This work | *C. jejuni* NCTC11168 WT CSS-0032 |
| *C. jejuni* NCTC11168 $\Delta cj1643$ | This work | *C. jejuni* PflC deletion CSS-4087 |
| *C. jejuni* NCTC11168 $\Delta cj0892c$ | This work | *C. jejuni* PflD deletion CSS-4081 |
| *C. jejuni* NCTC11168 Cj1643-3xFLAG | This work | *C. jejuni* PflC-3xFLAG CSS-4720 |
| *C. jejuni* NCTC11168 $\Delta cj0892c$ + *cj0892c-sfgfp* | This work | *C. jejuni* PflD-sfGFP CSS-4666 |
| *C. jejuni* NCTC11168 $\Delta cj0892c$ + *cj0892c-sfgfp*, PflA-3xFLAG | This work | *C. jejuni* PflD-sfGFP, PflA-3xFLAG CSS-5714 |
| *C. jejuni* NCTC11168 $\Delta cj0892c$ + *cj0892c-sfgfp*, PflB-3xFLAG | This work | *C. jejuni* PflD-sfGFP, PflB-3xFLAG CSS-5716 |
| *C. jejuni* 81-176 $\Delta flgQ$ / *pDRH7476* | This work | *C. jejuni* FlgQ-mCherry DRH7516 |
| *E. coli* DH5α | Lab stock | Cloning strain |
| *E. coli* BL21(DE3) | Lab stock | Protein expression strain |
| *C. jejuni* 81-176 *rpsL*<sup>Sm</sup> $\Delta pflA$ | Ref<sup>9</sup> | *C. jejuni* DAR1124 |
| *C. jejuni* 81-176 *rpsL* $\Delta pflA/pDAR3417$ | This work | *C. jejuni* DAR3447 |
| C. jejuni 81-176 *rpsL* $\Delta pflA/pDAR1604$ | This work | *C. jejuni* DAR3477 |
| C. jejuni 81-176 *rpsL*<sup>Sm</sup> $\Delta pflB$ | Ref<sup>9</sup> | *C. jejuni* DAR981 |
| C. jejuni 81-176 *rpsL* $\Delta pflB/pDAR3414$ | This work | *C. jejuni* DAR3451 |
| C. jejuni 81-176 *rpsL* $\Delta pflB/pDAR965$ | This work | *C. jejuni* DAR3479 |
| *C. jejuni* 81-176 $\Delta flaA flaB$<sup>S397</sup> *flhF*<sup>D321A</sup> $\Delta cheY$ | This work | CCW-locked *C. jejuni* with subpolar short filaments for bead assay |
| C. jejuni 81-176 *rpsL* $\Delta flgX/pDRH8743$ | This work | DRH8754 |
| Deposited data |  |  |
| Raw single particle analysis movies | This work | EMPIAR-11580 (DOI: 10.6019/EMPIAR-10016) |
| Whole motor map | This work | EMD-16723 |
| Lathed LP-ring focused refinement map | This work | EMD-16723 (additional volume) |
| Lathed C-ring focused refinement map | This work | EMD-16723 (additional volume) |
| Lathed MS-ring focused refinement map | This work | EMD-16723 (additional volume) |
| Focused periplasmic scaffold map | This work | EMD-16724 |
| $\Delta pflC$ subtomogram average | This work | EMD-17415 |
| $\Delta pflD$ subtomogram average | This work | EMD-17416 |
| *pflA*<sub>$\Delta 16-168$</sub> subtomogram average | This work | EMD-17417 |
| FlgQ-mCherry subtomogram average | This work | EMD-17419 |
| C-ring subtomogram average | This work | EMD-19642 |
| Recombinant DNA |  |  |
| Source of *cat* cassette for chloramphenicol resistance | Ref<sup>81</sup> | pRY109 |
| *E. coli*-*C. jejuni* shuttle vector | Ref<sup>81</sup> | pRY112 |
| pRY112 with 76-bp fragment containing *cat* promoter with start codon and in-frame BamHI restriction site cloned into the XbaI and XmaI sites | This work | pDAR1003 |
| pDAR1003 with DNA encoding *mcherry* and stop codon cloned in-frame with respect to the *cat* start codon and BamHI site into the XmaI and EcoRV sites | This work | pDAR1006 |
| Fusion with *flgQ* from codon 2 to the penultimate codon cloned into the BamHI site of pDAR1006 to create a FlgQ-mCherry fusion | This work | FlgQ-mCherry plasmid pDRH7476 |
| *E. coli*-*C. jejuni* shuttle vector | Ref<sup>81</sup> | pRY108 |
| *E. coli*-*C. jejuni* shuttle vector containing *cat* promoter and start codon for expression of genes for complementation | Ref<sup>82</sup> | pECO102 |
| *E. coli*-*C. jejuni* shuttle vector containing *cat* promoter and start codon followed by DNA encoding an in-frame N-terminal FLAG tag | Ref<sup>78</sup> | pDAR965 |
| pRY108 with 206 base pair fragment containing *flaA* promoter and start codon with in-frame SpeI site cloned into the XbaI and BamHI sites | This work | pDAR1425 |
| E. coli-C. jejuni shuttle vector containing flaA promoter and start codon followed by DNA encoding an in-frame N-terminal FLAG tag | Ref<sup>83</sup> | pDAR1604 |
| pECO102 with codon 2 to penultimate codon of pflB and an in-frame C-terminal FLAG epitope cloned into the BamHI site | This work | pDAR3414 |
| pDAR1425 with codon 2 to penultimate codon of pflA and an in-frame C-terminal FLAG epitope cloned into the BamHI site | This work | pDAR3417 |
| pECO102::*flgX-lysozyme* | This work | pDRH8743 |
| pLIC-PflA | This work | Full-length PflA (16-788) |
| pLIC-PflAα | This work | PflA TPR regions (169-788) |
| pLIC-PflA<sub>N</sub> | This work | PflA N-terminal half (16-454) |
| pLIC-PflB | This work | Soluble PflB (113-820) |
| pLIC-PflC | This work | Full-length PflC (17-364) |
| pLIC-PflC<sub>Δ236-349</sub> | This work | PflC C-terminal truncation (17-235) |
| pLIC (Amp<sup>R</sup>) | Franziska Sendker | Cloning vector backbone |

